# Structural dynamics of cytochrome P450 3A4 in the presence of substrates and cytochrome P450 reductase

**DOI:** 10.1101/2021.03.09.434557

**Authors:** Julie Ducharme, Irina F. Sevrioukova, Christopher J. Thibodeaux, Karine Auclair

## Abstract

Cytochrome P450 3A4 (CYP3A4) is the most important drug-metabolizing enzyme in humans and has been associated with harmful drug interactions. The activity of CYP3A4 is known to be modulated by several compounds, as well as by the electron transfer partner, cytochrome P450 reductase (CPR). The underlying mechanism of these effects however is poorly understood. We have used hydrogen-deuterium exchange mass spectroscopy (HDX-MS) to investigate the impact of CPR and three different substrates (7-benzyloxy-4-trifluoromethyl-coumarin, testosterone and progesterone) on the conformational dynamics of CYP3A4. Here, we report that interaction of CYP3A4 with substrates or with the oxidized or reduced form of CPR leads to a global rigidification of the CYP3A4 structure. This was evident from a suppression of deuterium exchange in several regions of CYP3A4, including those known to be involved in protein-protein interactions (C-helix) as well as substrate binding and specificity (B’-, E-helices and K/β1-loop). Furthermore, the bimodal isotopic distributions observed for some CYP3A4-derived peptides were drastically impacted by CPR and/or substrates, suggesting the existence of stable CYP3A4 conformational populations that are perturbed by ligand/CPR binding. The results have implications for understanding the mechanisms of allostery, ligand binding, and catalysis in CYP enzymes.

## INTRODUCTION

Cytochrome P450 enzymes (CYPs) are a large family of heme-containing monooxygenases that share a similar three-dimensional architecture. Most CYPs utilize O_2_ and electrons donated by a redox partner to catalyze substrate oxidation. Cytochrome P450 reductase (CPR) is the primary electron transfer partner of mammalian CYPs. Importantly, polymorphisms in the human *cpr* gene are associated with complex metabolic disorders, such as defects in steroid metabolism.^1^ It has been suggested that the *cpr* mutations may differentially affect the ability of the mutant CPR proteins to functionally interact with their endogenous CYP partners.^1^ CPR is a diflavin reductase that contains one molecule each of flavin adenine dinucleotide (FAD) and flavin mononucleotide (FMN). The flavin cofactors enable CPR to shuttle electrons from NADPH (an obligate 2e-donor) one at a time to CYP to support oxygen activation and, ultimately, substrate oxidation. During these electron transfer steps, CPR undergoes large conformational changes from an open to a closed form.^2–4,56^ In addition, various studies have reported that CPR may induce conformational changes in CYPs that promote inter-protein electron transfer.^6^ However, the current understanding of conformational changes induced by redox partners on CYPs is based mostly on the bacterial P450cam-putidaredoxin (Pdx) system.^7–9^

Human CYP3A4 is central to the metabolism of more than 50% of all pharmaceuticals, and is often implicated in drug interactions and chemo-resistance.^10,11^ This enzyme is well known to exhibit complex ligand-dependent allosteric and cooperative behaviors.^12^ Whereas many studies have focused on elucidating the allosteric effects of substrates,^13–16^ effectors,^14,17,18^ other CYPs,^19,20^ and the membrane,^21,22^ very little attention has been given to understanding how CPR affects CYP conformational dynamics. Since CYP-CPR interactions may be modulated by substrates, effectors and other CYP enzymes,^23–25^ a better understanding of the phenomenon will improve our ability to predict drug metabolism and interactions.

In the present study, we employ hydrogen-deuterium exchange mass spectrometry (HDX-MS) to characterize the conformational flexibility of CYP3A4 in the presence of CPR and/or its substrates 7-benzyloxy-4-trifluoromethyl-coumarin (BFC), testosterone (TST), or progesterone (PRG). The HDX-MS technique probes the dynamics of structural elements in proteins by monitoring the time-dependent exchange of deuterium atoms between the solvent and backbone amide groups.^26^ The rate of deuterium uptake is highly sensitive to the local hydrogen bonding environment and, as such, can be used to deduce ligand binding interactions and/or to infer the existence of conformational changes under different conditions (*e*.*g*. ligand bound/unbound states).^27^ A decrease in deuterium exchange typically reflects a structural organization, whereas an increase in deuterium uptake suggests enhanced flexibility.

As presented herein, our HDX-MS results reveal an overall increase in the rigidity of CYP3A4 upon interaction with CPR or substrates. Interestingly, the structural elements affected by CPR and substrates are largely shared, suggesting that binding of CYP3A4 to either CPR or substrate may stabilize the conformation poised for catalysis. This conclusion is supported by the coalescence of bimodal isotopic distributions of CYP3A4 peptides derived from the B’-, C-, E-helices and β1-sheet, suggesting a shift in the CYP3A4 conformational ensemble to a more homogeneous population of structures upon ligand/CPR binding. Finally, the dynamic structural elements of CYP3A4 perturbed by CPR are not restricted to the CYP-CPR interface, but also include regions associated with substrate binding and specificity. Cumulatively, these data build on the growing body of evidence for the functional relevance of conformational dynamics in CYP3A4.^28–30^

## METHODS

### Protein expression and purification

The *N-*terminally truncated Δ3-13 CYP3A4 was expressed and purified as previously described.^17^ Truncation of the transmembrane helix is commonly employed to improve protein expression yield and solubility and does not significantly impact enzymatic function under *in vitro* conditions.^31,32^ The protein was stored at -80 °C in 0.1 M potassium phosphate buffer (pH 7.4) containing 10% glycerol. Rat cytochrome P450 reductase (CPR) was expressed and purified as previously reported^33^ except for one important modification: the liquid growth medium was supplemented with a filter-sterilized, fully-dissolved riboflavin solution (0.2 mg/mL, 50:50 acetonitrile:H_2_O, pH 11), which was added at a volume ratio of 5 mL/L of the growth medium at the time of inoculation. Amino acid sequences of CYP3A4 and CPR are shown in **Figure S1**.

### Hydrogen-deuterium exchange protocol

Separately, the CYP3A4 and CPR solutions were concentrated to ca. 100 μM using Spin-X concentrators (30 kDa MWCO, 0.5 mL, Corning). All deuterium exchange reaction mixtures consisted of 1 μM CYP3A4 in deuterated potassium phosphate buffer (0.1 M KPi-D_2_O, pD 7.0) to ensure a final D_2_O concentration >98% (D_2_O/H_2_O), which is critical for maximizing deuterium uptake. As needed, CPR (4 μM), NADPH (1 mM), and/or substrate (100 μM of BFC, TST, or PRG) were added to the reaction mixture. Stock solutions were prepared for NADPH (60 mM in Milli-Q) and the substrates (20 mM in DMSO). Prior to the hydrogen-deuterium exchange reaction, when appropriate, the concentrated stocks of CYP3A4 and CPR were pre-incubated together for 20 minutes to allow for optimal interaction between the two proteins. The substrate stock solutions were pre-mixed in deuterated buffer before addition to the enzyme solution. HDX reactions were initiated by the addition of the enzyme(s) to deuterated buffer (with or without substrate). At the desired time points (0.5 - 60 min), aliquots of the reaction mixture (50 μL) were immediately quenched with pH 1.7 buffer (100 μL, 100 mM KPi) to yield a final pH of 2.5, thus ensuring minimal amide H/D back exchange. The quenched samples were immediately flash frozen in liquid nitrogen and stored at -80°C. Each reaction was performed in triplicate on the same day using the same stock solutions.

### MS data acquisition

All HDX-MS experiments were performed on a Waters Synapt G2-Si with HDX technology, and MS data acquisition was performed as previously reported.^34^ One at a time, the quenched reaction samples (150 μL, stored at -80°C in 1.5 mL Eppendorf tubes) were thawed for 70 s in a water bath at 32°C and then loaded into a 40 μL injection loop of the HDX manager. The samples were injected in the HDX manager exactly 2 min after they were removed from the freezer. In the HDX manager, the protein was cleaved by elution (100 μL/min flow rate of 0.1% formic acid in Milli-Q water) through an online immobilized pepsin column for 3 min at 15°C. The resulting peptic peptides were trapped on a C18 guard column held at 0.4°C before reverse-phase chromatographic separation using a Waters BEH C18 UPLC column (1 × 100 mm). Prior to loading peptides, the column was equilibrated in 97:3 (v/v) Milli-Q water:acetonitrile containing 0.1% formic acid. The peptides were eluted at 0.4°C with a linear gradient of 3-100% acetonitrile containing 0.1% formic acid applied over 10 min. The samples were ionized by electrospray ionization (ESI) with a capillary voltage of 2.8 kV, sampling cone of 30 V, source offset of 30 V, and desolvation temperature of 175°C. Data were collected in positive ion and resolution modes. The ionized peptides were then further separated in the gas phase by travelling wave ion mobility using a wave velocity of 650 m/s, wave height of 40 V, a bias voltage of 3 V, and a nitrogen pressure of 3.1 mbar. After exiting the ion mobility region, the peptides were subjected to two alternating collision energy regimes applied over 0.4 s intervals: a low collision energy of 6 V and a high collision energy ramp of 21 – 44 V. The high energy regime fragments ions by collision induced dissociation (CID), while the low energy regime conserves the peptide precursor masses. This process allows the matching of peptide fragments with their respective parent ions and enables peptide identification with high confidence. A [glu-1]-fibrinopeptide B (Glu-Fib) external lock mass standard was always run in parallel with all samples to enable correction of the peptide m/z values. An average variance in the determination of deuterium uptake values for this specific workflow was previously determined to be 0.087 Da with a standard deviation of 0.095 Da.^34^ These values are similar to the previously reported peptide-level continuous exchange, bottom-up HDX-MS deuterium uptake differences.^35,36^

### MS data processing

A peak list of the CYP3A4 peptides reproducibly detected by our workflow was generated. This list was determined by analyzing reference samples for CYP3A4 prepared in triplicate in protiated potassium phosphate buffer (100 mM, pH 2.5) and subjected to the HDX-MS workflow described above. The peptide reference lists were generated by uploading the raw MS data in the Protein Lynx Global Server (PLGS) software (Waters) as previously described.^34^ PLGS scans the raw data and associates the detected peptide peaks with expected masses. Spectral assignments were scored by a variety of user-defined criteria: ppm error, difference in chromatographic retention time, difference in ion mobility drift time, and number of MS/MS ion matches. The PLGS output was uploaded into DynamX 3.0 (Waters) for additional thresholding and data analysis. The final peptide list was restricted to include only peptides with m/z values within 5 ppm of the theoretical mass, that produced a minimum of 0.2 product (fragment) ions per amino acid, and that were detected in all three replicates. This resulted in a final list of 147 peptides covering 89.1% of the CYP3A4 sequence with an amino acid redundancy of 4.25.

### MS data analysis

DynamX 3.0 was used to quantify the level of deuterium uptake for each peptide under all conditions tested. DynamX 3.0 calculates a centroid mass from the isotopic distribution of deuterated peptide m/z signals. The centroid mass is then compared to the similarly determined centroid mass for the non-deuterated reference peptides. The deuterium uptake is quantified based on this comparison and is reported either as relative uptake (in Da) or relative fractional uptake (RFU; %), where the absolute deuterium uptake is normalized by the total number of amide protons in the peptide. Data was curated manually to ensure that all peptide assignments by the software were correct. Differential HDX was used to compare two states by subtracting the deuterium uptake data for each peptide for the two states of interest. We ensured that all compared peptides were from the same charge state, and that the reference and deuterated peptide had matching chromatographic retention times, ion mobility drift times, and m/z values. A summary of the conditions tested is provided in **Table S2**. Bimodal isotope distributions were fitted with HX-express v2 software.^37^

### Assessing HDX-MS variance

The overall reproducibility of the workflow was established as previously described.^34^ All difference data reported in this work represent a sum of the HDX difference values for each peptide at each of the analyzed time points. The significant deuterium uptake difference between compared states was determined using the Deuteros software package versions 1.0 or 2.0 developed by Politis and coworkers.^38^ The two versions of the software use different statistical tests to calculate the significance of deuterium uptake differences between the compared HDX profiles. The hybrid-test of Deuteros 2.0 was used for most states comparison.^39^ The less stringent statistical test of Deuteros 1.0 was only used to extract the significant differences between [CYP+BFC:CYP+TST/PRG] and [CYP+TST:CYP+PRG] (**Figure 5**).^38^ The RFU difference for the differential profiles of interest was mapped onto the CYP3A4 structure (PDB: 1W0F). The RFU normalizes the total level of deuterium uptake to the theoretical uptake maximum, based on the length of the peptide.

### Equilibrium titrations

Titration of CYP3A4 with TST and PRG was conducted at room temperature in 0.1 M potassium phosphate buffer, pH 7.4, supplemented with 10% glycerol. Substrates were dissolved in DMSO and added to a 1.5 μM protein solution in small aliquots, keeping the total volume of added titrant to < 2% of the final volume. After each addition, the reaction mixture was allowed to equilibrate for at least 20 min. UV-visible absorption spectra were recorded when the signal was stable. Titration plots were constructed from the difference spectra and fit to the Hill equation to derive dissociation constants (*K*_D_) and Hill coefficient (*n*_H_).

### BFC hydroxylation assay

The activity of CYP3A4 towards BFC was determined by measuring the initial rates of the enzyme-catalyzed debenzylation of BFC to yield 7***-***hydroxy***-***4-(trifluoromethyl)coumarin (HFC) using a microtiter plate reader with fluorescence detection. The assay was performed in potassium phosphate buffer (0.1 M, pH 7.4, 10% glycerol) containing CYP3A4 (0.5 μM), CPR (2 μM), BFC (100 μM from an initial stock of 30 mM in DMSO) and NADPH (1 mM from a 25 mM stock solution in deionized water) in a total reaction volume of 150 μL. The final DMSO concentration in the assay was below 1%. Prior to initiating the reaction, CYP3A4 and CPR were incubated in assay buffer for 1 h at room temperature to allow proper interaction. BFC was then added to the solution, before transferring to individual plate wells. The reaction was initiated by the addition of NADPH. HFC production was monitored for 2 h (Ex/Em: 410 nm/530 nm) at 37°C, and the signal measured was converted to μM/min using an HFC calibration curve. Initial rates were measured at different BFC concentrations (10-100 μM) and HFC formation was monitored for 90 min. Slopes were calculated from the initial linear portions of the kinetic curves. The kinetic data were fitted with both the Michaelis-Menten and Hill equations using GraphPad Prism 7.0. Better fits (higher *R*^2^ values) were obtained with the Hill equation.

### TST and PRG hydroxylation assays

The enzyme activity of CYP3A4 with the substrates TST and PRG in the presence of either CPR/NADPH or the cofactor surrogate cumene hydroperoxide (CHP) was monitored by LC-MS. The reaction mixtures contained CYP3A4 (0.5 μM), PRG or TST (10-150 μM from an initial stock of 1-15 mM in DMSO), CPR (2 μM) and NADPH (1 mM from a 25 mM stock in deionized water), or CHP (0.4 mM) in 0.1 M KPi buffer. When CPR was used, CPR and CYP3A4 were first pre-incubated together in KPi buffer at room temperature for 1 h, and the reactions were initiated with the addition of NADPH and allowed to proceed at 37°C for a total 75 minutes. When CHP was used, the reactions were initiated with CHP and the total reaction time was 6 minutes. All reactions were performed in duplicate. For each substrate concentration, reaction rates were obtained from the linear portion of the kinetic curve containing three time points (CHP: 2, 4, 6 min; CPR: 45, 60, 75 min). At the appropriate time point, 150 ml of the reaction mixture was quenched by addition of 0.5 mL DCM. The substrates and products were extracted in DCM (2 × 0.5 mL) and the combined organic layers were dried under N_2_. The dried samples were re-dissolved in acetonitrile (0.1 mL) before LC-UV-MS analysis. The samples were eluted on a C-18 Omega Polar HPLC column (Phenomenex, 250 × 4.6 mm, 5 mm particle size) using 50:50 acetonitrile:H_2_O for 3 min, followed by a linear increase to 80% acetonitrile over 9 min, and another linear increase to 95% acetonitrile over 8 min. The products were detected by their absorption at 244 nm and their corresponding masses were confirmed by MS detection in positive ion mode. Hydroxylated TST products eluted between 7-10 min and TST eluted at 15 min. Hydroxylated PRG products eluted between 12-16 min and PRG eluted at 21 min. Calibration curves of TST and PRG were used to convert the absorption signals of the product peaks into concentration (μM).

## RESULTS

### Overview of the HDX workflow

To investigate the structural dynamics of CYP3A4, a continuous-labeling, bottom-up HDX-MS approach was utilized (**Table S2**). Deuterium exchange on the backbone amide groups of CYP3A4 was achieved by exposing the protein to a deuterated buffer for various periods of time ranging from 0.5 to 60 min, in the presence or absence of CPR (oxidized or reduced), BFC, TST or PRG. Our workflow resulted in the reproducible detection of 57-64 peptides, providing a CYP3A4 sequence coverage of 74-80%. Regions with no coverage included the A’’-helix, small portions of the D- and I-helices, and the F’- and G-helices. The statistical test used to determine the significance of the changes observed in deuterium uptake (DU) across the CYP3A4 structure was the hybrid test as implemented in Deuteros version 2.0,^39^ with confidence intervals varying between 99-99.9%. This hybrid statistical test combines individual significance testing with a globally estimated threshold calculated from the variance of all data points.^40^ All significant peptides detected in our study are detailed in **Figures S3** and **S4**, and comparisons between results obtained under different conditions can be visualized in butterfly plots (**Figure S5**). The significant uptake changes over time were summed in order to capture more subtle changes in HDX.

CYP3A4 is known to oligomerize in solution.^41,42^ Thus, we carried out a control experiment to ensure that CYP3A4 aggregation was not interfering with the deuterium uptake observed under our conditions. Since the level of protein aggregation is proportional to concentration, we performed an HDX-MS experiment in which we varied the CYP3A4 concentration by 10-fold.^28^ The results show no significant change in deuterium uptake across the protein structure (74% sequence coverage) for all four CYP3A4 concentrations tested (0.5, 1, 2 and 5 mM).

### Impact of the binding of oxidized CPR on the structural dynamics of CYP3A4

We first studied the structural dynamics of CYP3A4 in the presence of oxidized CPR (*ox*CPR) for comparison to the free enzyme ([CYP:CYP+*ox*CPR]). The oxidation state of CPR was determined spectroscopically (**Figure S7**). A 4-fold excess of CPR to CYP3A4 was used in our experiments, as this ratio was determined to be optimal for enzymatic activity towards the substrates BFC, TST and PRG (**Figure S6**). Out of 64 CYP3A4 peptic peptides compared with and without *ox*CPR, 26 peptides showed a significant difference (CI 99.9%) in deuterium uptake (**Figures S3, S8**). The respective sum of RFU differences for these peptides at all D_2_O exposure times (0.5, 5, 30 min) is mapped onto the CYP3A4 structure in **Figure 1**. Interestingly, structural dynamics changes associated with *ox*CPR binding were distributed throughout the CYP3A4 structure rather than localized at the putative CPR-CYP interface (includes the C-helix of CYP3A4). Furthermore, binding of *ox*CPR results in an overall decrease in deuterium uptake, which suggests a global reduction in the flexibility of CYP3A4. Peptide 319-333 located in the J-helix is a notable exception that becomes more flexible in the presence of *ox*CPR (Δuptake = +7.7 Da, 61% RFU). As expected, a substantial deuterium uptake difference was observed at the binding interface between CYP3A4 and CPR (peptide 126-137, C-helix), with a total decrease in uptake of -3.2 Da (−33% RFU) upon CPR binding. Large decreases in deuterium uptake were also observed for the B’- and E-helices (− 5.5 Da and -3.4 Da, respectively) and the β1/K-loop (−3.0 Da, -61% RFU) upon *ox*CPR binding. These regions are associated with substrate binding and specificity, suggesting that interaction of CYP3A4 with CPR may help to define substrate binding sites. Interestingly, the proximal loop involved in the heme redox potential regulation (peptide 429-441) also becomes more rigid.

**Figure 1.**
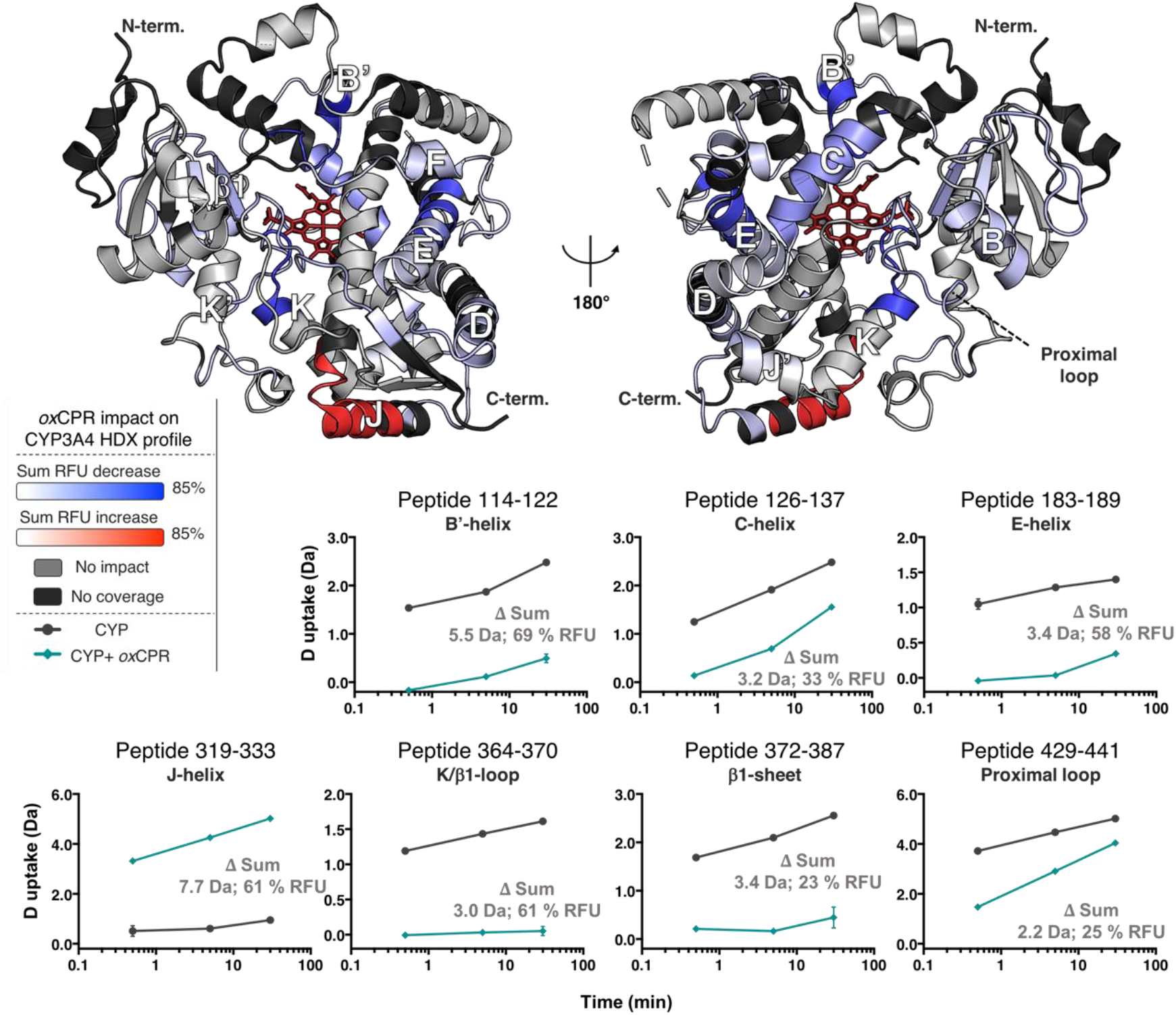
Impact of *ox*CPR binding on the HDX profile of CYP3A4. The differential HDX profile of [CYP:CYP+*ox*CPR] is shown. The significant sum RFU difference (CI 99.9%) of three time points (0.5, 5, 30 min) is mapped onto the CYP3A4 structure (PDB: 1W0F), which is shown in both distal (left) and proximal (right) views. Blue is used to display regions that undergo a decrease in deuterium uptake upon interaction with *ox*CPR, whereas red is used to designate regions that undergo an increase in deuterium uptake over time. The light gray color shows regions unaffected by substrate binding, and the regions shown in black were not covered in the MS analysis. Deuterium uptake time courses are shown for the most impacted peptides in the bottom panels. The sum DU and RFU difference is shown for each (CI 99.9%). The plots for CYP3A4 are in black and those for the CYP3A4+*ox*CPR complex are colored in teal.

### Impact of the binding of reduced CPR on the structural dynamics of CYP3A4

CPR is believed to undergo a drastic conformational change upon reduction.^2–4,56^ The reduced form of CPR (*red*CPR) was also reported to have a higher affinity for CYPs than *ox*CPR.^43^ Therefore, we hypothesized that binding of *red*CPR might affect the structural dynamics of CYP3A4 differently than *ox*CPR. To investigate this, we generated a differential deuterium exchange profile comparing free CYP3A4 with the CYP3A4-*red*CPR complex ([CYP:CYP+*red*CPR]). Reduction of CPR was achieved by the addition of 1 mM NADPH and was confirmed by UV/Vis absorption spectroscopy (**Figure S7**). The deuterium exposure times (0.5, 5, 30 min) were selected to ensure that CPR remained reduced across all time points in the HDX reaction (**Figure S7**). The full differential HDX profile of [CYP:CYP+*red*CPR] is shown in **Figure 2**, and the complete list of significant peptides is provided in **Figure S3**. All deuterium uptake plots containing significant peptides are reported in **Figure S10**.

**Figure 2.**
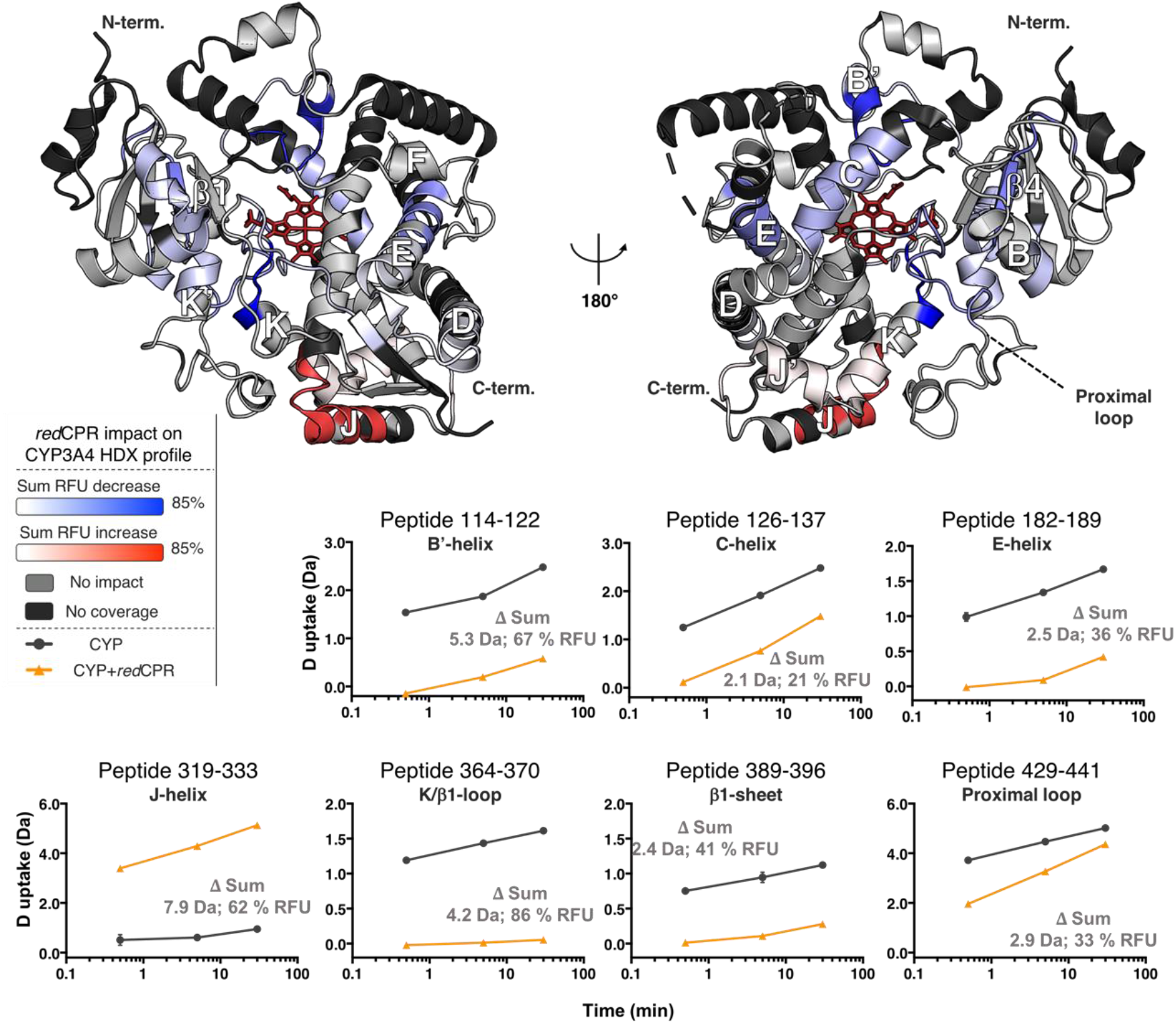
Impact of *red*CPR binding on the HDX profile of CYP3A4. The differential HDX profile of [CYP:CYP+*red*CPR] is shown. The significant sum RFU difference (CI 99.9%) of three time points (0.5, 5, 30 min) is mapped onto the CYP3A4 structure (PDB: 1W0F), which is shown in both distal (left) and proximal (right) views. Blue is used to display regions undergoing a decrease in deuterium uptake upon interaction with *red*CPR, whereas red is used to designate regions showing an increase in deuterium uptake over time. The light gray color shows regions unaffected by substrate binding, and the regions shown in black were not covered in the MS analysis. Deuterium uptake plots for the most important changes observed are shown at the bottom. The sum DU and RFU difference is shown for each (CI 99.9%). The plots for CYP3A4 are in black and those for the CYP3A4 + *red*CPR mixture are colored in orange.

Overall, the effect of *red*CPR binding on the conformational dynamics of CYP3A4 was found to be very similar to that of *ox*CPR binding, and was characterized by an obvious rigidification of the CYP3A4 structure relative to the free enzyme. The specific comparison of the [CYP+*ox*CPR:CYP+*red*CPR] states, however, revealed seven significantly different peptides (CI 99%), many of which were slightly more flexible in the [CYP:CYP+*red*CPR] system than in the [CYP:CYP+*ox*CPR] system (**Figure 3)**. These regions include the D/E-loop (0.4 Da, 3% RFU), the J/J’-loop (0.6 Da, 9% RFU) and the H/I-loop (0.6 Da, 4% RFU). Interestingly, the proximal loop, which bears the heme ligating Cys422, directly influences the heme redox potential, also becomes significantly more dynamic in the presence of *red*CPR (0.5 Da, 6% RFU) compared to *ox*CPR. Moreover, the middle portion of the I-helix (peptide 307-314), which contains the conserved Thr309 residue critical for proton relay during oxygen activation, is more flexible in the [CYP+*red*CPR] state. The only region found to undergo a decrease in deuterium uptake in the [CYP+*red*CPR] state relative to the [CYP+*ox*CPR] state was the K’-helix (−0.6 Da, 11% RFU). The potential relevance of this K’-helix rigidification is not immediately clear.

**Figure 3.**
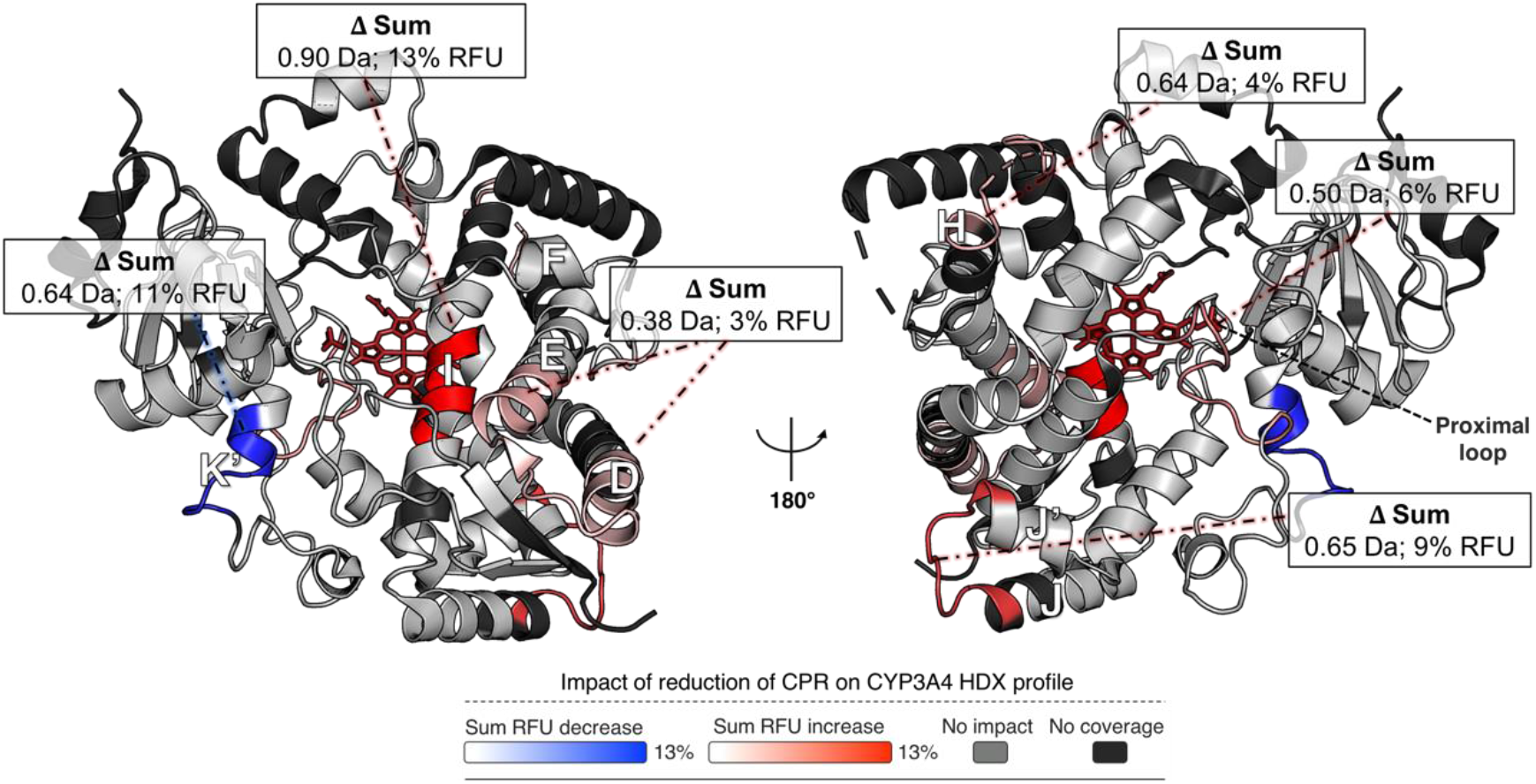
Impact of the binding of *red*CPR compared to that of *ox*CPR on the HDX profile of CYP3A4. The differential HDX profile comparing [CYP+*ox*CPR:CYP+*red*CPR] is shown. The sum RFU difference of three time points (0.5, 5, 30 min) is mapped onto the CYP3A4 structure (PDB: 1W0F), which is shown in both distal (left) and proximal (right) views. The blue color reflects a decrease in deuterium uptake upon reduction of CPR, whereas the red color designates regions undergoing an increase in deuterium uptake over time. Overall, the CYP3A4 structure is more dynamic in the presence of *red*CPR. The light gray color identifies regions unaffected by the oxidation state of CPR, and the regions shown in black were not covered in the MS analysis. The sum DU and RFU difference is shown for all statistically significant changes (CI 99%).

### Impact of substrate binding on the structural dynamics of CYP3A4

We next looked at the structural dynamics of CYP3A4 in complex with each of three different substrates. The differential HDX profiles of CYP3A4 in the presence/absence of BFC, TST and PRG ([CYP:CYP+BFC/TST/PRG]) were obtained for two D_2_O exposure time points (5, 60 min), which revealed 16, 16 and 12 peptides, respectively, for which deuterium exchange was differentially affected by substrate binding (CI 99%). The substrate concentration (100 μM) was selected to be higher than the respective dissociation constants (by 2-to-5 fold) without exceeding the substrate solubility limit in phosphate buffer. Overall, the differential HDX profiles acquired with each of the three substrates were very similar and comparable to the profile obtained with CPR binding, *i*.*e*. characterized by a general rigidification of the CYP3A4 structure (**Figure S9**). The CYP3A4 structural dynamic changes shared by all three enzyme-substrate complexes are shown in **Figure 4**. The CYP3A4 regions most affected by substrate binding include the B’-, E- and K’-helices. Interestingly, the C-helix, involved in CPR binding, also becomes more rigid upon substrate binding. These data suggest that CPR and substrates may function synergistically on the dynamic structure of CYP3A4 to modulate each other’s binding affinities.

**Figure 4.**
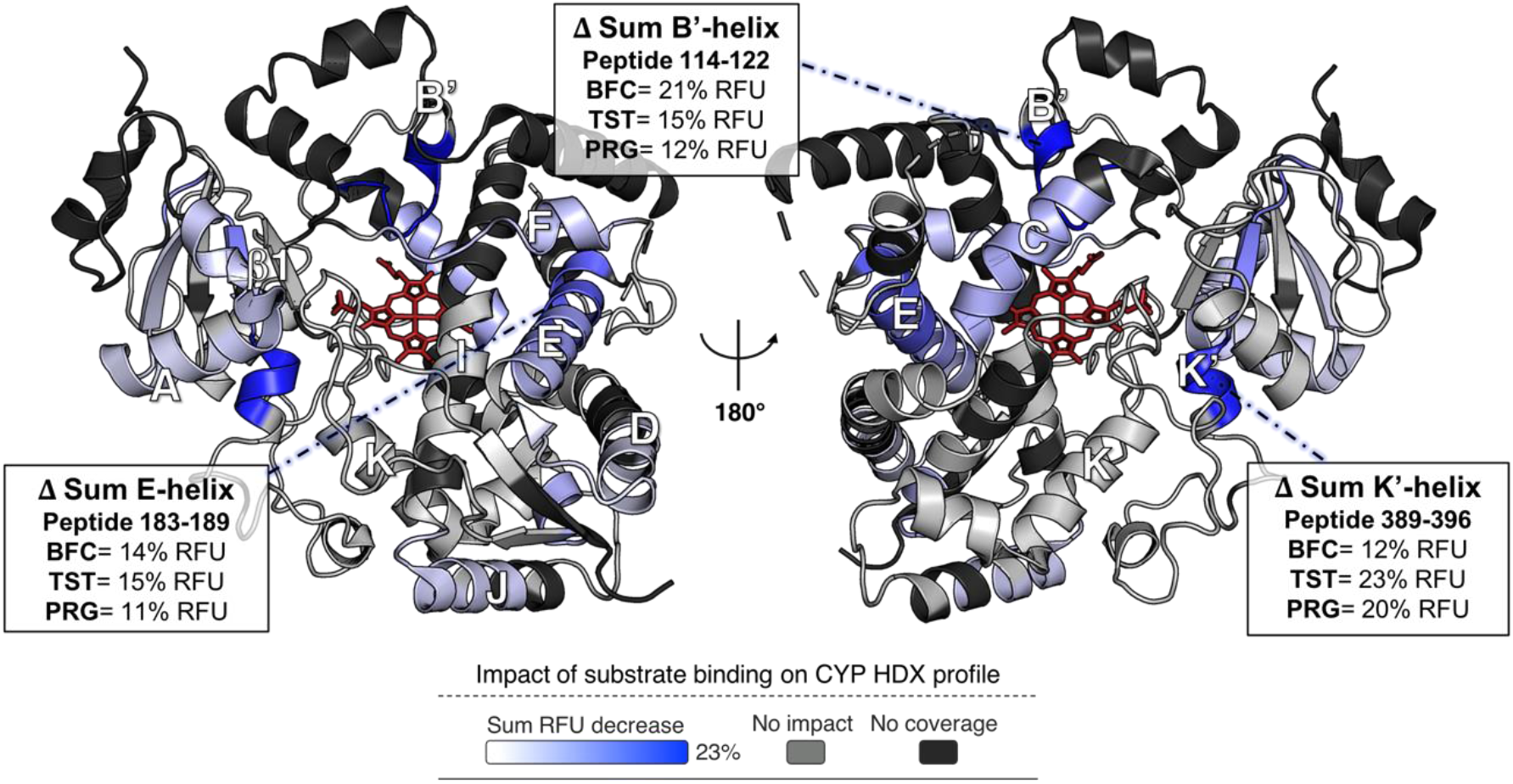
Substrate binding triggers a general rigidification in CYP3A4 that mirrors the effect of CPR binding. Differential HDX profile of [CYP:CYP+BFC/TST/PRG] ([S] = 100 μM). The sum RFU difference of two time points (5, 60 min) is mapped onto the CYP3A4 structure (PDB: 1W0F), which is shown in both distal (left) and proximal (right) views. The blue color signifies a decrease in deuterium uptake in the presence of substrate and is interpreted as a general organization/rigidification of the protein structure. The light gray color indicates regions that were unaffected by substrate binding and the black denotes regions that were not covered in the MS analysis. The sum RFU difference is shown for the three most affected regions: the B’-helix (peptide 114-122), the E-helix (peptide 183-189) and the K’-helix (peptide 389-396).

**Figure 5.**
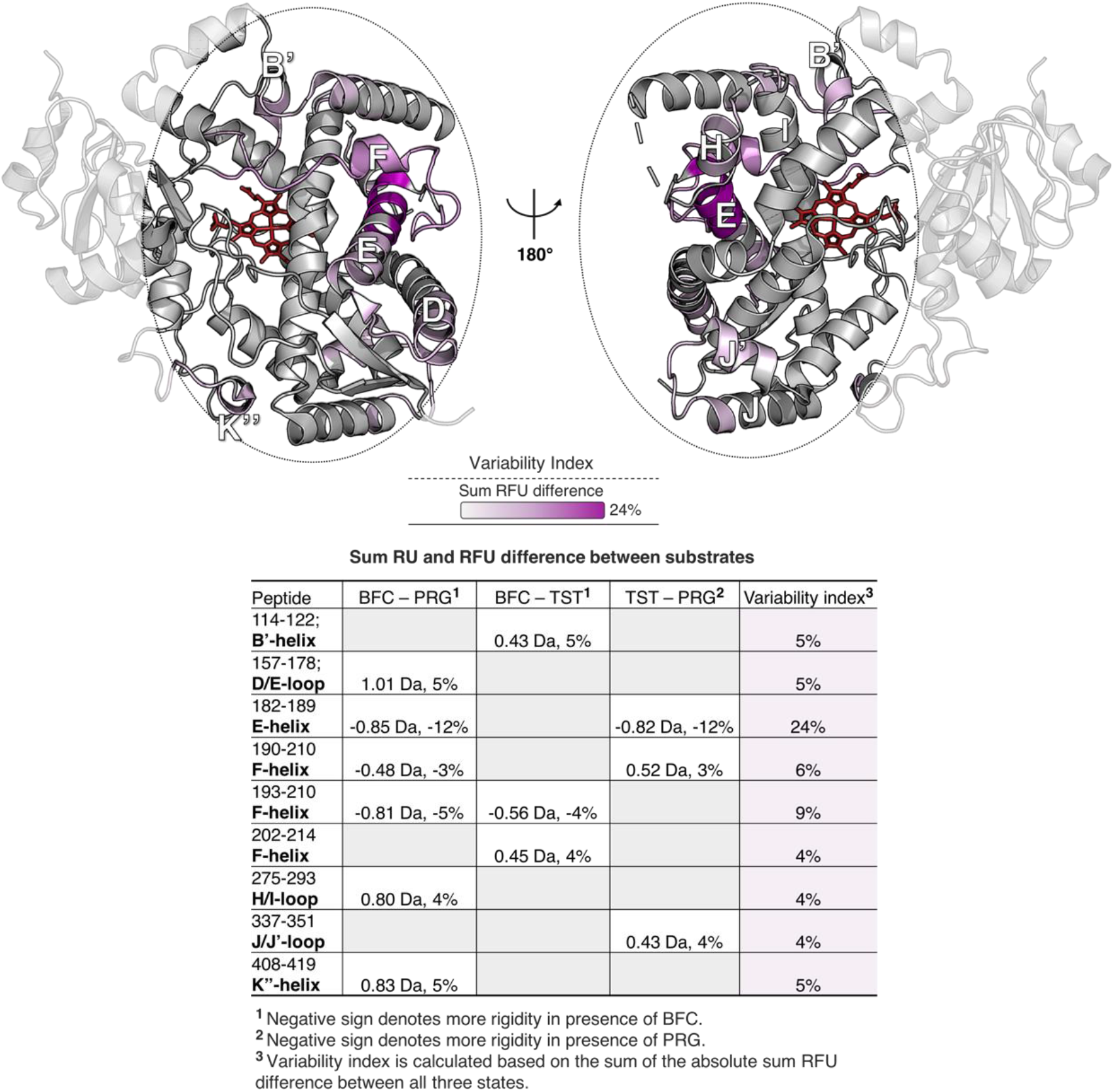
Variability in the structural dynamics of CYP3A4 upon binding to BFC, TST or PRG. The regions showing significant variability (CI 99%) upon binding of the investigated substrates are mapped on the structure of CYP3A4 (PDB: 1W0F) in both distal (left) and proximal (right) views. The variability index is calculated based on the absolute sum RFU differences obtained for each substrate as detailed in the table. In general, these structural elements that exhibit variable deuterium uptake cluster together in the three-dimensional structure of CYP3A4.

Despite the high degree of similarity in the overall structural dynamic changes induced by substrate binding, several regions of CYP3A4 exhibited subtle differences in uptake between the various substrates (**Figure 5**). These regions included the end portion of the E-helix (peptide 182-189) and the F-helix region (peptide 193-210), which exhibited the highest variation in deuterium uptake between substrates, as well as the D/E-loop, B’- and K’-helices. Although these regions are widely separated on the polypeptide chain, most of them are in close proximity in the three-dimensional structure, suggesting that the conformational dynamics in these CYP3A4 elements may be coupled to one another. These subtle differences in the dynamics of substrate-bound CYP3A4 may reflect heterogeneous binding modes of the different substrates, illustrating the power of HDX-MS to reveal slight structural differences between biochemical states of an enzyme.

Comparison of the relative impact of CPR and substrate binding on deuterium uptake by CYP3A4 reveals a more extensive rigidification induced by CPR binding (**Figure 6**). Most regions rigidified by CPR binding are also rigidified by substrate binding, including the B’-, C-, D-, E-, F-helices and the β1-sheet region. The only structural motif of CYP3A4 that exhibited opposite changes in deuterium uptake was the J-helix, which became more flexible in the presence of CPR but slightly more rigid upon substrate binding. The J-helix is directly connected to the I-helix and may impact the position of the I-helix relative to the heme. Movement of the I-helix is required to make space for productive substrate binding.

**Figure 6.**
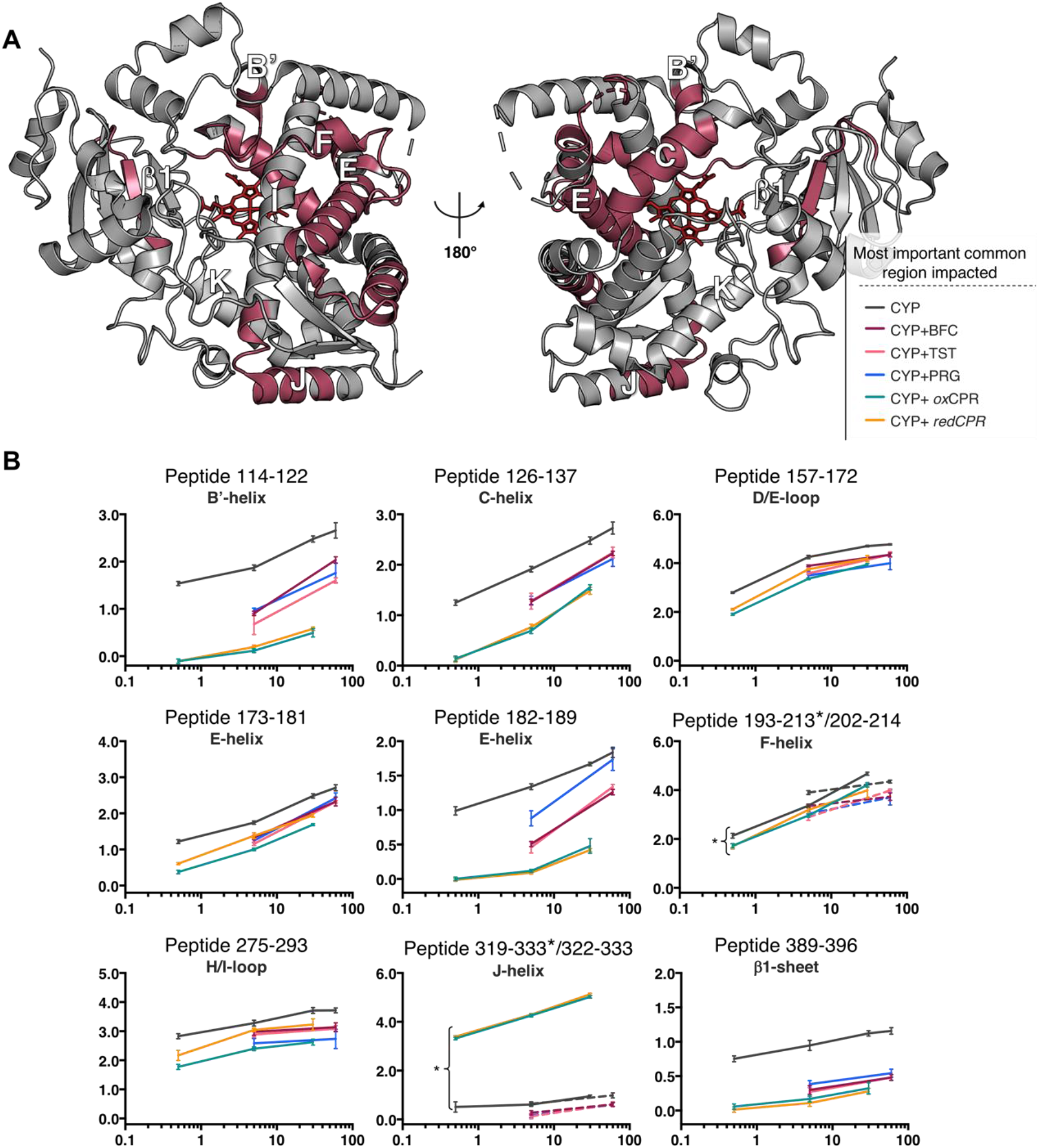
CYP3A4 peptides that exhibited significant deuterium uptake differences upon binding to either CPR or substrate. **A**) The location of peptides is indicated in raspberry on the CYP3A4 structure (PDB: 1W0F) showing distal (left) and proximal (right) views. **B**) Deuterium uptake plots of these common regions showing uptake data for all conditions tested in this study. CYP uptake plots are shown in black, CYP+BFC in purple, CYP+TST in pink, CYP+PRG in blue, CYP+*ox*CPR in teal and CYP+*red*CPR in orange. Slightly different coverages were obtained for the F- and J-helices in the CPR- and substrate-bound states. In these cases, two different peptides are displayed on the deuterium uptake plots. Peptides 193-213 and 319-333 are plotted as solid lines and designated with an asterisk. Peptides 202-214 and 322-333 are plotted as doted lines.

### Impact of CPR and substrate binding on bimodal isotopic distribution

Under native conditions, most proteins uptake deuterium in the EX2 kinetic regime, where the protein samples unfolded conformations that refold at rates that are higher than the rate of deuterium exchange. This results in peptide mass spectra with a unimodal isotope distribution.^27^ Careful analysis of the mass spectrum of each peptic peptide revealed a few spectra that deviated from this common EX2 behavior, showing instead a bimodal isotope pattern characteristic of the EX1 kinetic regime. Such an isotope distribution results from peptides that sample long-lived open conformational states that fully exchange with solvent deuterium before refolding into the closed state. A total of five peptides from free CYP3A4 showed a clear bimodal distribution pattern (**Figure 7**). These include peptides derived from the B’- and C-helices, the end-portion of the E-helix, the F-helix and the β1-sheet region. With the exception of peptide 374-387 (in the β1-sheet), these peptides show a significant decrease in deuterium uptake upon CPR or substrate binding (**Figures 1, 2, 4, S3-S5**). Interestingly, the uptake difference of peptide 374-387 in the β1-sheet did not significantly change upon CPR/substrate binding, but the bimodal distribution was almost completely gone in the CYP-CPR and CYP-subtrates complexes. This suggests that ligand/CPR binding induces a more rapid conformational equilibrium in the β1-sheet that exchanges in the EX2 regime. In all cases, except for the F-helix region, a large decrease in the relative abundance of the highly exchanged population of the free enzyme was observed upon ligand binding, consistent with the general rigidification of the enzyme upon CPR or substrate binding. The effect was more pronounced upon CPR binding. Ligand/CPR binding apparently shifts the equilibrium of CYP3A4 conformations towards a more structured form of the enzyme that samples the open (rapidly exchanging) conformation less often. The interaction of CPR with CYP3A4 almost completely suppressed the highly-exchanged population of the B’-, C-, E-helices and β1-sheet regions. In contrast, substrate binding decreased the abundance of the highly exchanged population of the C-helix (believed to directly interact with CPR) and the β1-sheet (located on the distal face near the heme group), but did not significantly affect the bimodal isotope distribution of the E-helix peptide. Interestingly, the bimodal distribution of the B’-helix (which is involved in substrate binding and specificity) seemed to be more sensitive to the nature of the substrate. No drastic changes are observed in the bimodal distribution of the F-helix, except that in all cases the slowly exchanged populations appear to uptake less deuterium. Again, this data illustrates the ability of HDX-MS to reveal subtle changes in structural dynamics induced by different ligand binding modes.

**Figure 7.**
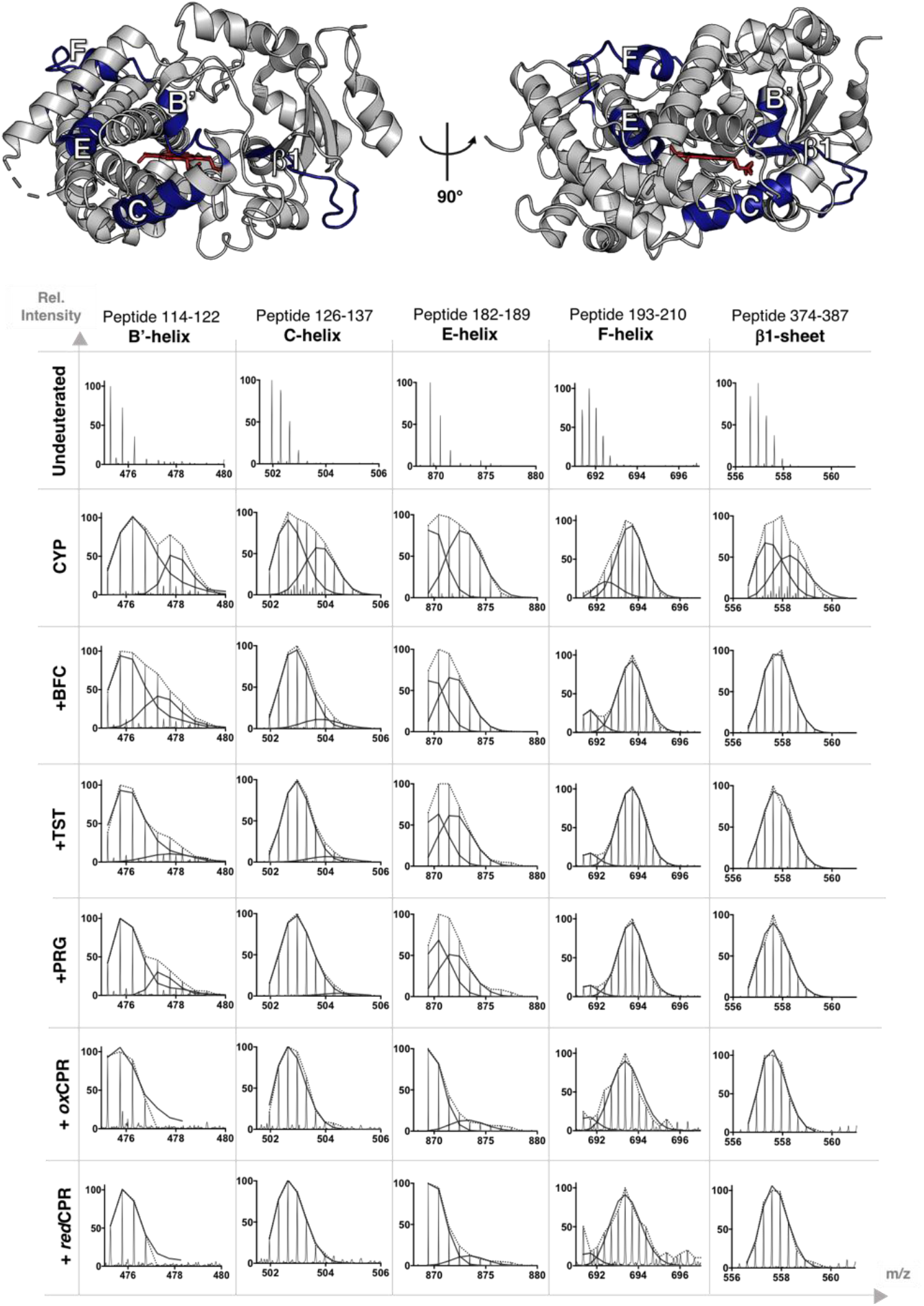
Mass spectra for the five HDX peptides of free CYP3A4 that displayed a bimodal isotope distribution. The B’-, C-, E-, F-helices and the β1-sheet region are mapped onto the CYP3A4 structure (1W0F) in deep blue. The mass spectra of these peptides are compared for CYP3A4 alone or in the presence of *ox*CPR, *red*CPR, BFC, TST, or PRG. The data is presented for the highest D_2_O exposure time point (60 min for substrates, 30 min for CPR). The dotted lines indicate the peptide mass envelope, and the solid lines represent the fitted deconvolution of the isotope distribution to a sum of two binomial populations.

### Impact of CPR on functional activity and substrate binding affinity of CYP3A4

To relate the HDX-MS data to functional activity and substrate binding, we conducted enzymatic and equilibrium titrations assays. Kinetics of BFC, TST and PRG oxidation was measured using either CPR (4–fold excess) and NADPH, or the hydrogen peroxide donor cumene hydroperoxide (CHP) as a surrogate of the redox partner (**Figure 8**). CHP is known to sustain the activity of CYP3A4 and bypass the electron transfer steps normally catalyzed by CPR in the native system.^44^ Both CPR/NADPH and CHP concentrations were optimized to achieve maximum catalytic activity (**Figure S6**). The results revealed that the functional positive cooperativity (sigmoidal behavior) for these substrates^45^ was generally more pronounced in the presence of CPR, which suggests a direct contribution of CPR to this phenomenon. In agreement with our previous results,^44^ the enzyme activity with all substrates was significantly higher with CHP than with CPR/NADPH, likely as a result of bypassing the rate-limiting electron transfer steps. Due to poor solubility of steroids in aqueous solutions and the low catalytic turnover of the CPR-supported system, *V*_*max*_ for TST hydroxylation could not be reached, precluding an accurate kinetic characterization of this system. The faster rates achieved with CHP, however, allowed us to calculate steady state kinetic constants. At least for BFC and PRG, the data are consistent with a smaller functional cooperativity with CHP than with CPR. Spectrophotometric equilibrium titrations were next performed to evaluate dissociation constants (*K*_d_). Whereas PRG and TST induce a type I spectral shift (blue shift) in the λ_max_ of the Soret band of CYP3A4, binding of BFC is not associated with a significant perturbation in the heme absorption. Therefore, the titrations were only conducted with PRG and TST. To prevent substrate turnover, *ox*CPR was used. We found that the presence of *ox*CPR slightly decreased the extent of the PRG-induced spectral shift (**Figure 9A**) and reduced the dissociation constant (*K*_d_) for PRG by nearly 2-fold, from 35 to 19 μM (**Figure 9C**). In contrast, *ox*CPR enhanced the TST-dependent spectral shift (**Figure 9B)**, and slightly weakened the binding affinity for TST from 46 to 55 μM (**Figure 9D**). The Hill coefficients (*n*_H_) for binding of both substrates were only modestly affected by *ox*CPR (**Figure 9C, D**). Thus, *ox*CPR binding to CYP3A4 has a distinct effect on the binding of different substrates.

**Figure 8.**
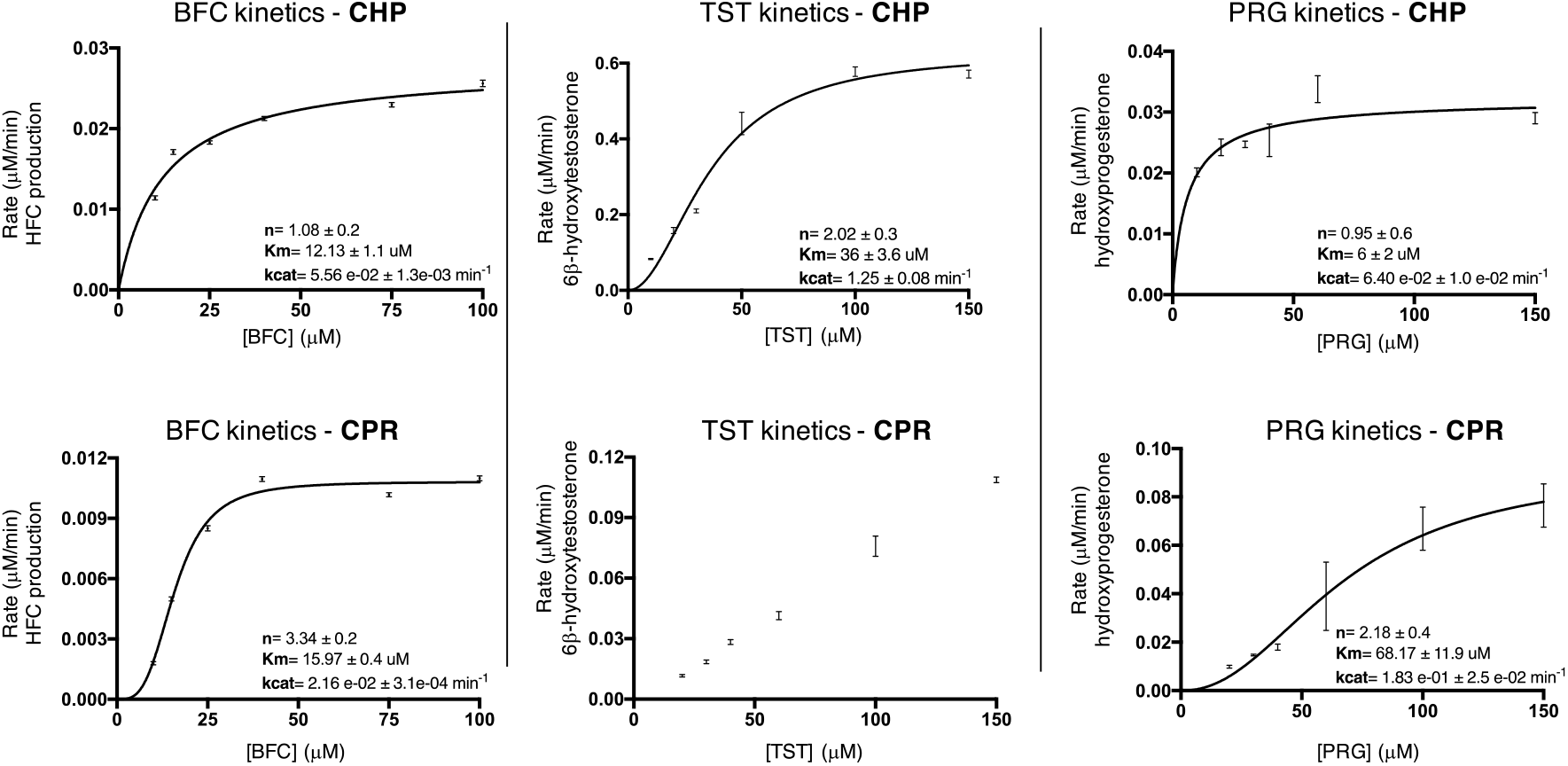
Impact of CPR on the functional activity of CYP3A4. Kinetics of BFC, TST or PRG oxidation by CYP3A4 (0.5 μM) was measured in the presence of CPR (2 μM) or CHP (0.4-0.6 mM). Error bars represent standard deviations of duplicate or triplicate measurements.

**Figure 9.**
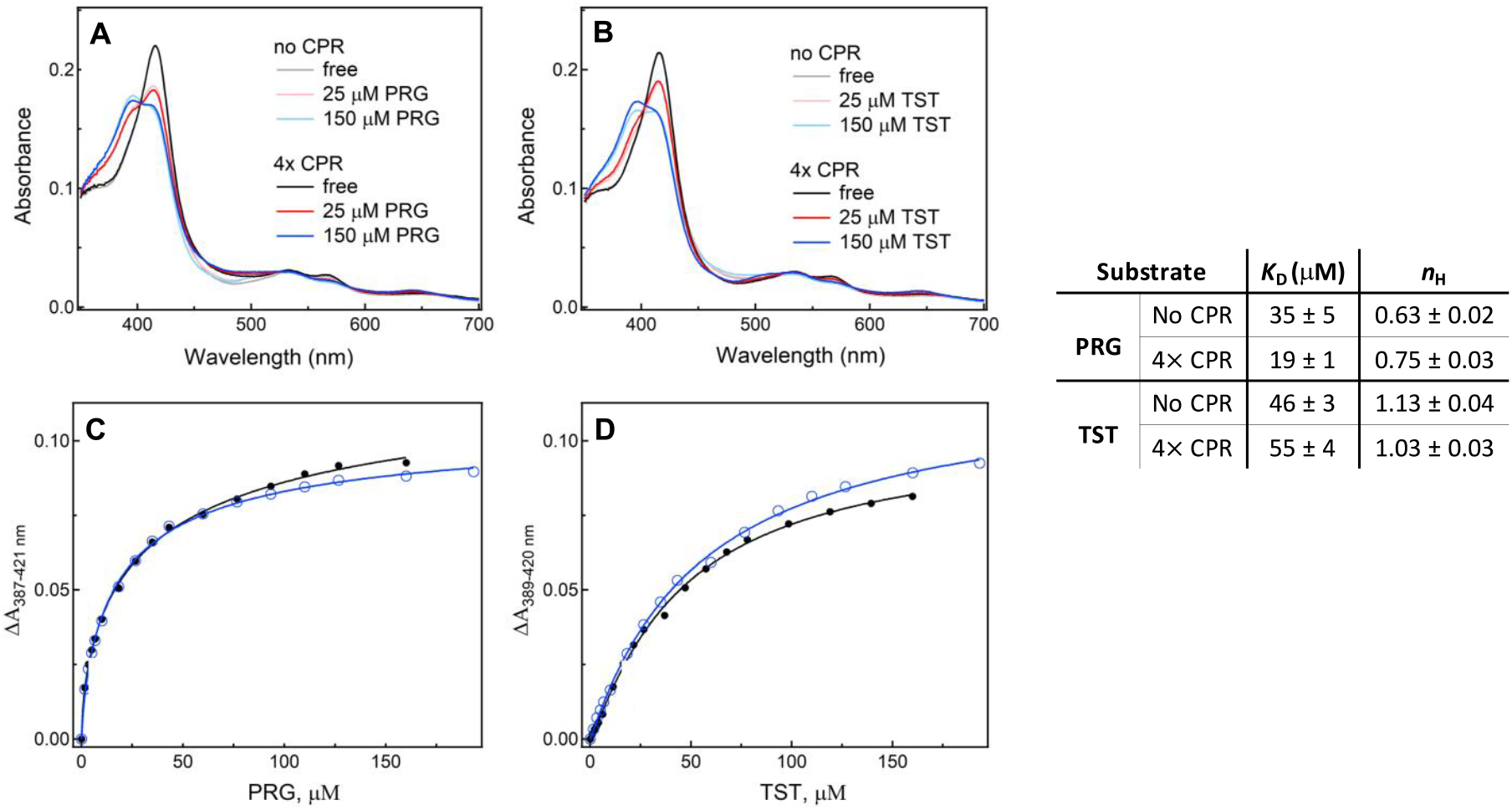
The impact of *ox*CPR on the spectral properties of CYP3A4 and its binding to substrate. **A** and **B**, absorption spectra of CYP3A4 (1.5 μM) in the presence of variable concentrations of PRG/TST and a four-fold molar excess *ox*CPR (6 μM). Substrate addition shifts the heme absorption from 421 nm to 387 nm, indicative of the low-to-high spin transition in the heme iron. **C** and **D**, titration plots for PRG and TST, respectively, fitted with the Hill equation. The *K*_d_ and *n*_H_ values derived from the fits are shown in the table.

Together, the titration and kinetic data suggest that CPR modulates both the binding properties and the functional activity of CYP3A4. Interestingly, the dynamics of the proximal K’’/L-loop (bearing the cysteine ligated to the heme) becomes significantly more rigid in the presence of CPR. These changes in structural dynamics could potentially affect the heme redox potential and Soret absorption. Additionally, the impacts of substrates and CPR on the structural dynamics of CYP3A4 may be additive and could represent a structural basis for the modulation of CYP3A4 substrate binding and functional properties by CPR.

## DISCUSSION

In the present study, we investigated the effects of CPR and substrate binding on the conformational dynamics of human CYP3A4 – an enzyme that exhibits remarkably relaxed substrate specificity and complex allosteric behavior. Our approach utilizes hydrogen-deuterium exchange mass spectrometry (HDX-MS) to provide peptide-level, localized information on protein conformational dynamics. Human CPR is encoded by a single gene and serves as a redox partner to support the activity of various enzymes, including many CYPs and heme oxygenases. CYPs share a common fold and likely possess a similar CPR binding surface. It is well accepted that the negatively charged surface of the FMN-binding domain of CPR interacts with the positively charged concave surface on the proximal face of CYPs.^46^ In particular, mutagenesis and bioconjugation studies support a role for the basic residues of the C-helix of human CYP2B1, CYP2B4 and CYP2B6 in CPR binding.^47–49^ Both the deuterium uptake levels and isotope distribution data presented here are consistent with a direct interaction between the C-helix of CYP3A4 and CPR. Namely, rigidification of the C-helix (decreased deuterium uptake) supports the involvement of this CYP element in protein-protein interactions with CPR. Suppression of the highly exchanged EX1 population (as indicated by the bimodal mass distribution of peptides from the C-helix) can be interpreted as a perturbation in the CYP3A4 conformational ensemble towards a more structured form upon CPR binding.

Crystal structures of CPR have revealed important conformational changes upon flavin reduction.^2,4,50^ Molecular dynamics simulations and NMR studies also support the ability of CPR to sample both open and closed conformations.^5,51–55^ The closed form enables intra-protein electron transfer between the two flavin coenzymes, whereas the open form of CPR mediates inter-protein electron transfer to CYPs.^5^ This implies that the CYP-CPR complex may dissociate between the two distinct single electron transfer steps required for oxygen activation.^5,46^ Our HDX-MS results reveal a significant overall rigidification of CYP3A4 upon CPR binding, regardless of the oxidation state of the reductase. These data suggest that both *ox*CPR and *red*CPR may bind to similar CYP3A4 conformations and/or induce comparable conformational perturbation to CYP3A4 upon interaction. While the overall HDX profiles of the [CYP+*ox*CPR] and [CYP+*red*CPR] complexes are quite similar, reduction of CPR leads to an increase in the flexibility of the D/E-loop, the proximal K’’/L-loop, and the I-helix relative to the *ox*CPR state. The K’’/L-loop has been implicated in the regulation of the heme redox potential, whereas the I-helix bears the conserved Thr309 residue that mediates proton transfers during dioxygen activation.^32^ Conformational flexibility in the I-helix was also observed in P450cam and thought to be important for oxygen activation and binding of the redox partner Pdx.^8,56–58^ These data illustrate the potential for CPR binding to modulate the structural dynamics of CYP3A4 with a finely-tuned regioselectivity that depends on the redox state of the reductase.

Structural evidence is accumulating to support a mechanism by which redox partner binding may cause conformational changes in the CYP active site and even at the distal surface of the protein.^46^ This was first explored with P450cam, whose association with the electron donor (Pdx) was found to cause a shift from a closed to an open conformation of the P450 enzyme via motions of the F- and G-helices.^8,9^ A putative communication pathway has been proposed for the Pdx-induced conformational change via coupled motions of the B’-, C-, I-, D- and F-helices,^56^ which is consistent with our results on the CYP3A4-CPR pair (**Figures 1** and **S8**). Among all regions of CYP3A4 impacted by CPR binding, the structural rigidification was strongest in the B’-helix, which defines two major substrate access channels, and the active site K/β1-loop, which participates in substrate binding. A moderate change in dynamic properties of the F-helix was also observed. This region is important for achieving both the open and closed conformations of P450cam and CYP3A4,^9,59^ and is also implicated in substrate access and binding affinity. Together with our spectral and enzyme activity results (**Figures 8** and **9**), the HDX data suggests that association with CPR alters the ligand binding properties at the active site of CYP3A4.^60–62^

All three substrates tested were found to similarly influence the structural dynamics of CYP3A4 (**Figures 5** and **6**). This is of particular interest as it was suggested earlier that ligand binding to CYPs could induce conformational changes that favor/disfavor redox partner association.^41–45^ For example, Zhang et al.^7^ showed that binding of the substrate androstenedione decreased the affinity of human aromatase for CPR, and that the presence of CPR increased the rate of substrate binding. Our HDX-MS data suggest that both substrate and CPR binding may be shifting the CYP3A4 conformational ensemble to a structurally similar active state. Thus, both substrate and CPR may synergistically affect each other’s binding to CYP3A4 which, in turn, could contribute to the well-known positive kinetic cooperativity of this system.

Previous HDX-MS investigations with truncated (Δ3-12) CYP3A4 revealed small, but significant changes in deuterium uptake when the lipid nanodisc-incorporated and detergent-solubilized enzymes were compared.^30^ Overall, the deuterium uptake properties were similar, except for the dynamics of the F- and G-helices, which are expected to interact directly with the membrane mimic. These regions were found to be more rigid in the presence of the membrane nanodiscs. More recently, the same group reported the effect of midazolam binding to CYP3A4 in lipid nanodiscs.^29^ Under the conditions used (60 μM midazolam, 13.5 min D_2_O exposure time), the largest impact of ligand binding was observed for the F- and G-helices (increased flexibility) and the K/β1-loop (rigidification).^29^ This is consistent with our HDX results, as the same regions were impacted by BFC, TST or PRG. However, we observed rigidification of the F-helix region, which contrasts to the increased flexibility for this structural element reported by Atkins and co-workers. This difference may arise from the distinct experimental settings, *e*.*g*. lipid-free vs. membrane environment and/or utilization of distinct substrates. Nevertheless, our study provides the basis for future investigations on the membrane incorporated [CYP3A4+CPR] complex which could help to resolve some of these discrepancies.

In summary, this is the first HDX-MS study to examine the effects of CPR binding on the structural dynamics of CYP3A4, and provides new information on redox partner interactions and substrate binding in CYP enzymes. Our data suggest that (i) both *ox*CPR and *red*CPR trigger rigidification of CYP3A4 to a similar extent, most notably in the B’-, C-, E-helices and the K/β1-loop region, and (ii) CPR modulates the substrate binding properties and catalytic activity of CYP3A4 in a substrate dependent manner. Furthermore, strong similarities between the regions undergoing conformational changes upon redox partner association to CYP3A4 and P450cam were noted and may suggest a common allosteric mechanism for dioxygen binding/activation stemming from the redox partner binding. All three investigated substrates (BFC, TST and PRG) triggerred similar perturbations in CYP3A4 dynamics, but with some distinctions that may reflect different ligand binding modes. Together, our results help to better understand structural dynamics of CYP3A4 and its potential role in modulating catalytic function.

## Acknowledgments

Julie Ducharme was supported by scholarships from the CGCC and the FRQNT. We would like to thank Dr. J. R. Halpert for providing the CYP3A4 plasmid and Dr. C. B. Kasper for the CPR plasmid. We are also grateful for the generous support provided by Dr. M. Guttman for using the HX-express v2 software.

## Supporting Information

Amino acid sequences of human CYP3A4 and CPR, HDX experimental conditions, detailed differential HDX uptake data, CPR- and CHP-dependent activity plots for CYP3A4, and spectral confirmation of CPR reduction by NADPH.

## Accession Codes

Cytochrome P450 3A4 (CYP3A4): P08684.

## Notes

The authors declare no competing financial interest.

