## Supplemental Figures and Tables for "Structural dynamics of cytochrome P450 3A4 in the presence of substrates and cytochrome P450 reductase"

**Human CYP3A4 (Δ3-13)**
MALLLAVFLVLLYLYGTHSHGLFKKLGIPGPTPLPFLGNILSYHKGFCMFDMECHKKYGKVWGFYDGQQPVLAITDPDMIKTVLVKECYSVFTNR
RPFGPVGFMKSAISIAEDEEWKRLRSLLSPTFTSGKLKEMVPIIAQYGDVLVRNLRREAETGKPVTLKDVFGAYSMDVITSTSFGVNIDSLNNPQ
DPFVENTKKLLRFDFLDPFFLSITVFPFLIPILEVLNICVFPREVTNFLRKSVKRMKESRLEDTQKHRVDFLQLMIDSQNSKETESHKALSDLELVA
QSIIFIFAGYETTSSVLSFIMYELATHPDVQQKLQEEIDAVLPNKAPPTYDTVLQMEYLDMVVNETLRLFPIAMRLERVCKKDVEINGMFIPKGVVV
MIPSYALHRDPKYWTEPEKFLPERFSKKNKDNIDPYIYTPFGSGPRNCIGMRFALMNMKLALIRVLQNFSFKPCKETQIPLKLSLGGLLQPEKPVV
LKVESRDGTVSGAHHHH

**Rat CPR**
MGDSHEDTSATMPEAVAEEVSLFSTTDMVLFSLIVGVLTYWFIFRKKKEEIPEFSKIQTTAPPVKESSFVEKMKKTGRNIIVFYGSQTGTAEEFAN
RLSKDAHRYGMRGMSADPEEYDLADLSSLPEIDKSLVVFCMATYGEGDPTDNAQDFYDWLQETDVDLTGVKFAVFGLGNKTYEHFNAMGKYV
DQRLEQLGAQRIFELGLGDDDGNLEEDFITWREQFWPAVCEFFGVEATGEESSIRQYELVVHEDMDVAKVYTGEMGRLKSYENQKPPFDAKN
PFLAAVTANRKLNQGTERHLMHLELDISDSKIRYESGDHVAVYPANDSALVNQIGEILGADLDVIMSLNNLDEESNKKHPFPCPTTYRTALTYYLD
ITNPPRTNVLYELAQYASEPSEQEHLHKMASSSGEGKELYLSWVVEARRHILAILQDYPSLRPPIDHLCELLPRLQARYYSIASSSKVHPNSVHIC
AVAVEYEAKSGRVNKGVATSWLRAKEPAGENGGRALVPMFVRKSQFRLPFKSTTPVIMVGPGTGIAPFMGFIQERAWLREQGKEVGETLLYY
GCRRSDEDYLYREELARFHKDGALTQLNVAFSREQAHKVYVQHLLKRDREHLWKLIHEGGAHIYVCGDARNMAKDVQNTFYDIVAEFGPMEH
TQAVDYVKKLMTKGRYSLDVWS

**Figure S1.** Amino acid sequence of human CYP3A4 and rat CPR.

| Data Set | CYP3A4 | CYP3A4 + rCPR*ox* | CYP3A4 + rCPR*red* |
|---|---|---|---|
| HDX reaction details | 0.1 M KPi, pD = 7.0, RT, 1 μM CYP | 0.1 M KPi, pD = 7.0, RT, 1 μM CYP, 4μM CPR | 0.1 M KPi, pD = 7.0, RT, 1 μM CYP, 4μM CPR, 1mM NADPH |
| HDX time course (min) | 0.5, 5, 30, 60 | 0.5, 5, 30 | 0.5, 5, 30 |
| # of Peptides | 64 | 64 | 63 |
| Sequence coverage (%) | 77.7 | 79.9 | 75.3 |
| Redundancy | 1.95 | 1.88 | 1.93 |
| Replicates | 3 technical | 3 technical | 3 technical |
| Repeatability (maximum SD between replicates, Da) | 0.39 | 0.22 | 0.52 |

| Data Set | CYP3A4 + BFC | CYP3A4 + TST | CYP3A4 + PRG |
|---|---|---|---|
| HDX reaction details | 0.1 M KPi, pD = 7.0, RT, 1 μM CYP, 100 μM BFC | 0.1 M KPi, pD = 7.0, RT, 1 μM CYP, 100 μM TST | 0.1 M KPi, pD = 7.0, RT, 1 μM CYP, 100 μM PRG |
| HDX time course (min) | 5, 60 | 5, 60 | 5, 60 |
| # of Peptides | 57 | 58 | 58 |
| Sequence coverage (%) | 73.6 | 73.6 | 73.6 |
| Redundancy | 1.86 | 1.89 | 1.89 |
| Replicates | 3 technical | 3 technical | 3 technical |
| Repeatability (maximum SD between replicates, Da) | 0.62 | 0.59 | 0.58 |

**Table S2.** Summary tables of HDX-MS results for all experimental conditions tested. All experiments were performed under the same conditions, but the number of $D_2O$ exposure time points varied. The number and nature of the detected peptides varied slightly, but the percent sequence coverage remained higher than 76% for all states. Redundancy is a measure of the average number of peptides that cover each amino acid.

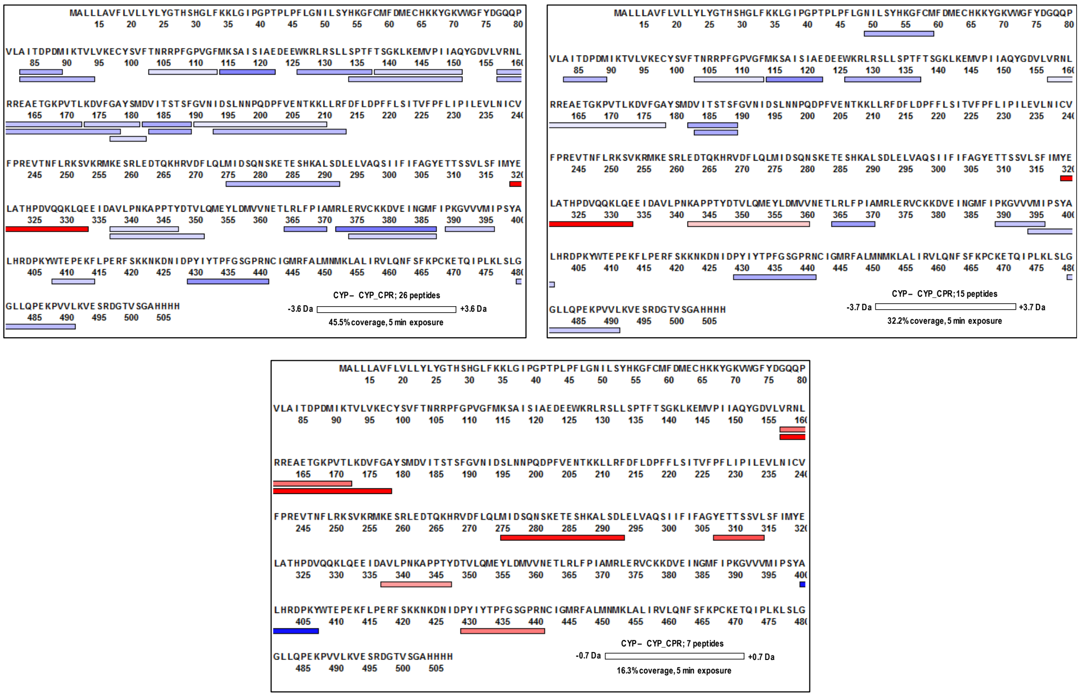


| Deuterium uptake difference: CYP – CYP + CPR*ox* | | | |
|---|---|---|---|
| Start | End | Sum Da | Sum RFU |
| 83 | 89 | 0.82 | 16.75 |
| 83 | 94 | 1.42 | 14.51 |
| 103 | 113 | 0.69 | 8.79 |
| 114 | 122 | 5.45 | 69.50 |
| 126 | 137 | 3.25 | 33.17 |
| 134 | 151 | 0.79 | 5.35 |
| 138 | 151 | 0.61 | 5.20 |
| 157 | 172 | 2.55 | 18.62 |
| 157 | 178 | 1.20 | 6.10 |
| 173 | 181 | 0.84 | 10.65 |
| 177 | 182 | 0.74 | 15.06 |
| 182 | 189 | 1.23 | 17.89 |
| 183 | 189 | 3.39 | 57.69 |
| 190 | 210 | 0.86 | 4.89 |
| 193 | 213 | 2.35 | 13.29 |
| 275 | 292 | 1.23 | 7.39 |
| 275 | 293 | 2.16 | 12.25 |
| 319 | 333 | -7.78 | -60.65 |
| 337 | 347 | 1.10 | 16.08 |
| 337 | 351 | 0.61 | 5.64 |
| 364 | 370 | 3.00 | 61.32 |
| 372 | 387 | 3.41 | 23.21 |
| 374 | 387 | 2.08 | 16.33 |
| 389 | 396 | 0.69 | 11.70 |
| 408 | 414 | 0.72 | 14.68 |
| 429 | 441 | 2.25 | 25.52 |
| 480 | 491 | 1.89 | 21.43 |

| Deuterium uptake difference CYP - CYP + CPR*red* | | | |
|---|---|---|---|
| Start | End | Sum Da | Sum %RFU |
| 49 | 59 | 1.52 | 15.48 |
| 83 | 89 | 0.76 | 15.58 |
| 103 | 113 | 0.75 | 9.51 |
| 114 | 122 | 5.26 | 67.05 |
| 126 | 137 | 2.11 | 21.55 |
| 157 | 178 | 0.98 | 4.98 |
| 182 | 189 | 2.50 | 36.51 |
| 183 | 189 | 1.00 | 16.94 |
| 319 | 333 | -7.88 | -61.84 |
| 342 | 360 | -0.77 | -4.94 |
| 364 | 370 | 4.19 | 85.57 |
| 389 | 396 | 2.41 | 40.95 |
| 394 | 401 | 0.82 | 13.93 |
| 429 | 441 | 2.95 | 33.45 |
| 480 | 491 | 1.05 | 11.87 |

| Deuterium uptake difference CYP + CPR*ox* – CYP + CPR*red* | | | |
|---|---|---|---|
| Start | End | Sum Da | Sum RFU |
| 157 | 172 | 0.39 | 2.82 |
| 157 | 178 | 0.68 | 3.49 |
| 275 | 293 | 0.64 | 3.63 |
| 307 | 314 | 0.90 | 13.12 |
| 337 | 347 | 0.65 | 9.46 |
| 400 | 407 | -0.65 | -11.00 |
| 429 | 441 | 0.50 | 5.62 |

**Figure S3.** Coverage map and tables of CYP3A4-derived peptides that undergo a significant change in deuterium uptake in the presence of *ox*CPR or *red*CPR. Coverage map bars and table entries shown in blue represent peptides that become more rigid in the presence of *ox*CPR or *red*CPR (i.e. less deuterium uptake), whereas peptides shaded in red become more flexible under the same conditions (i.e. more deuterium uptake).

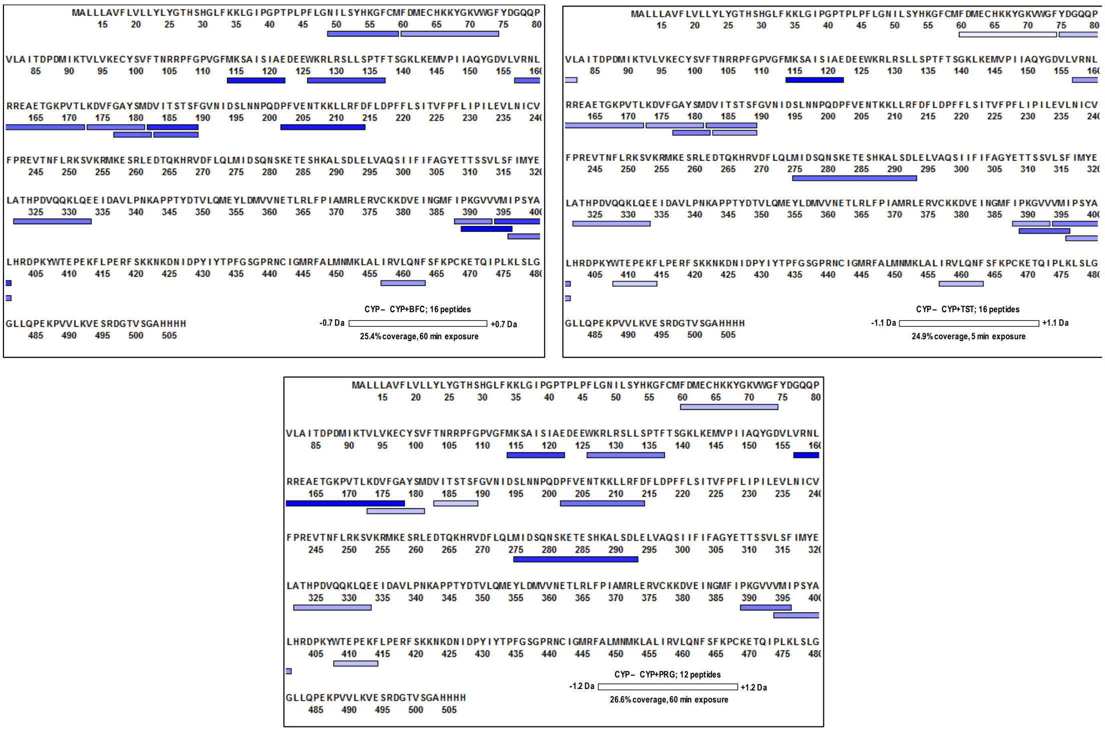


| Deuterium uptake difference: CYP - CYP_BFC | | | |
|---|---|---|---|
| Start | End | Sum Da | Sum %RFU |
| 49 | 59 | 0.40 | 4.07 |
| 60 | 74 | 0.40 | 2.94 |
| 114 | 122 | 1.67 | 21.34 |
| 126 | 137 | 0.65 | 6.60 |
| 157 | 172 | 0.36 | 2.63 |
| 173 | 181 | 0.42 | 5.38 |
| 177 | 182 | 0.41 | 8.47 |
| 182 | 189 | 0.84 | 12.21 |
| 183 | 189 | 0.82 | 13.97 |
| 202 | 214 | 0.56 | 5.19 |
| 322 | 333 | 0.37 | 3.76 |
| 388 | 393 | 0.38 | 9.63 |
| 389 | 396 | 0.68 | 11.59 |
| 394 | 401 | 0.66 | 11.22 |
| 396 | 401 | 0.78 | 20.02 |
| 457 | 463 | 0.39 | 6.60 |

| Deuterium uptake difference: CYP - CYP_TST | | | |
|---|---|---|---|
| Start | End | Sum Da | Sum %RFU |
| 60 | 74 | 0.52 | 3.83 |
| 75 | 82 | 0.41 | 7.05 |
| 114 | 122 | 1.14 | 14.61 |
| 157 | 172 | 0.67 | 4.86 |
| 173 | 181 | 0.60 | 7.71 |
| 177 | 182 | 0.42 | 8.62 |
| 182 | 189 | 0.91 | 13.29 |
| 183 | 189 | 0.86 | 14.66 |
| 202 | 214 | 1.04 | 9.67 |
| 275 | 293 | 0.63 | 3.55 |
| 322 | 333 | 0,53 | 5.40 |
| 388 | 393 | 0.76 | 19.50 |
| 389 | 396 | 1.37 | 23.30 |
| 394 | 401 | 0.68 | 11.64 |
| 396 | 401 | 0.41 | 10.39 |
| 408 | 414 | 0.41 | 8.35 |
| 457 | 463 | 0.42 | 7.09 |

| Deuterium uptake difference: CYP - CYP_PRG | | | |
|---|---|---|---|
| Start | End | Sum Da | Sum %RFU |
| 60 | 74 | 0.48 | 3.48 |
| 114 | 122 | 0.92 | 11.69 |
| 126 | 137 | 0.63 | 6.38 |
| 157 | 178 | 1.15 | 5.87 |
| 173 | 181 | 0.50 | 6.37 |
| 183 | 189 | 0.66 | 11.15 |
| 202 | 214 | 0.88 | 8.13 |
| 275 | 293 | 0.72 | 4.09 |
| 322 | 333 | 0.48 | 4.92 |
| 389 | 396 | 1.17 | 19.97 |
| 394 | 401 | 0.63 | 10.64 |
| 408 | 414 | 0.49 | 9.93 |

**Figure S4.** Coverage map and tables of CYP3A4-derived peptides that undergo a significant change in deuterium in the presence of either BFC, testosterone (TST), or progesterone (PRG). Coverage map bars and table entries highlighted in blue represent peptides becoming more rigid in the presence of substrates.

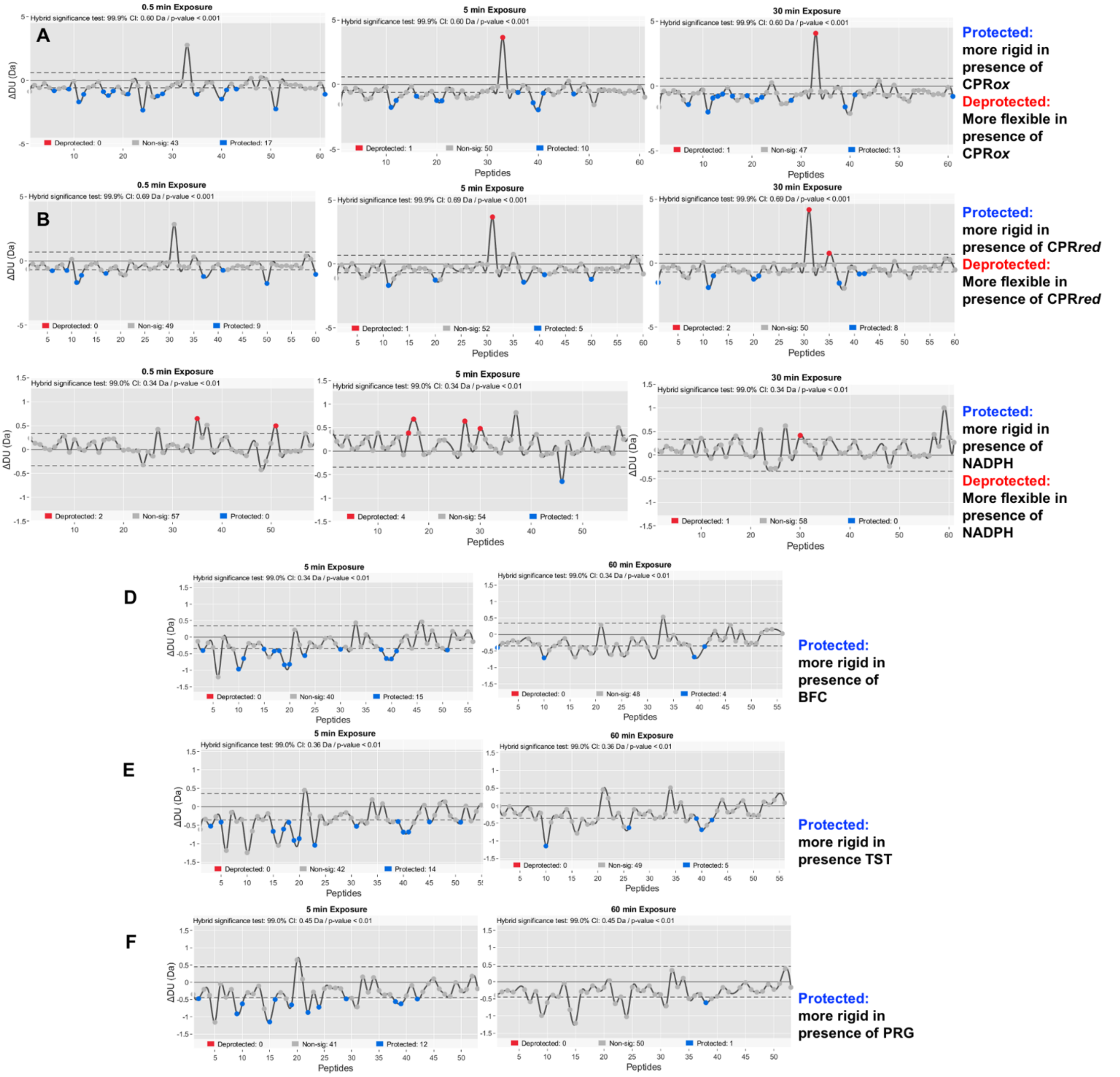


**Figure S5.** Butterfly plots mapping the significant differences among peptides: **A**) [CYP:CYP+*ox*CPR] (CI 99.9%); **B**) [CYP:CYP+*red*CPR] (CI 99.9%); **C**) [CYP+*ox*CPR:CYP+*red*CPR] (CI 99%); **D**) [CYP:CYP+BFC]; **E**) [CYP:CYP+TST]; **F**) [CYP+CYP+PRG]. Each dot represents a peptide and, when the dot is colored, it becomes considerably more rigid or more flexible in the presence of added CPR or substrate. For each set, the color coding is explained on the right. The y axis represents the uptake difference measured in Da.

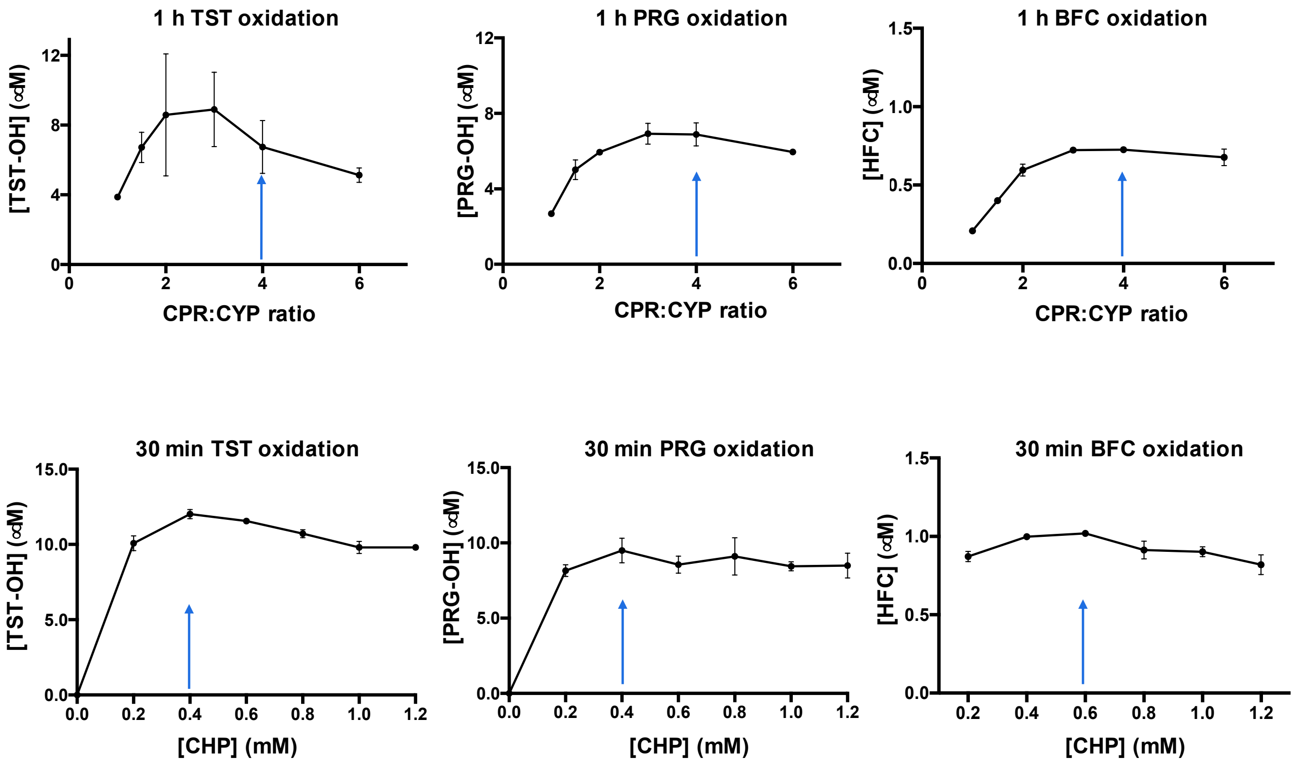


**Figure S6.** Concentration-dependent effect of CPR and CHP on the activity of CYP3A4 towards testosterone (TST), progesterone (PRG) and BFC.

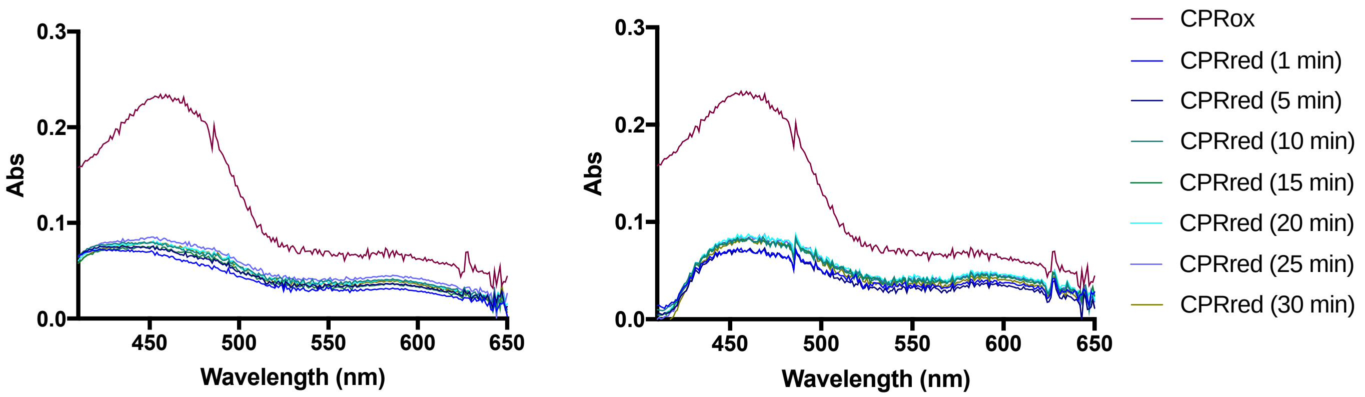


**Figure S7.** Absorption spectra of *ox*CPR and *red*CPR. The purple trace represents the spectrum of CPR in its oxidized form. Other CPR spectra were recorded at different time points (1-30 min) after addition of 1 mM NADPH. The reduction of CPR was measured both in the absence (left) and presence (right) of CYP3A4 (ratio 4:1 CPR to CYP3A4). Conditions: CYP3A4 (2.9 μM), CPR (11.7 μM), NADPH (1 mM), potassium phosphate buffer (0.1 M, pH 7.4, 10% glycerol) at room temperature.

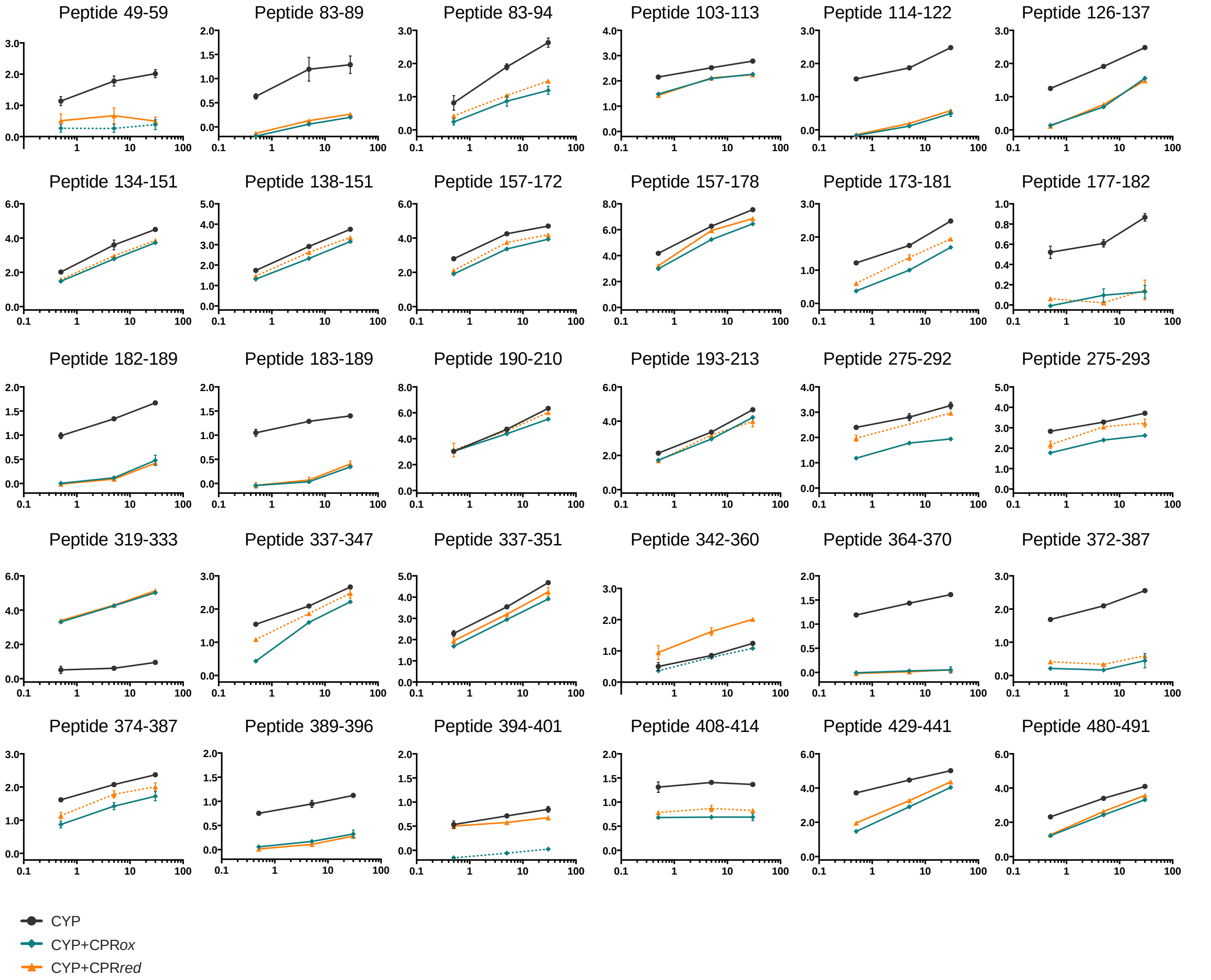


**Figure S8.** Deuterium uptake plots for all regions showing significant changes in the presence of either *ox*CPR or *red*CPR. The y-axis represents the deuterium uptake (Da) and the x-axis is time in a logarithmic scale. Black curves are for CYP alone, teal curves for CYP+*ox*CPR, and orange curves for CYP+*red*CPR. Dashed curves indicate that the difference in deuterium uptake was not significant (CI 99.9%).

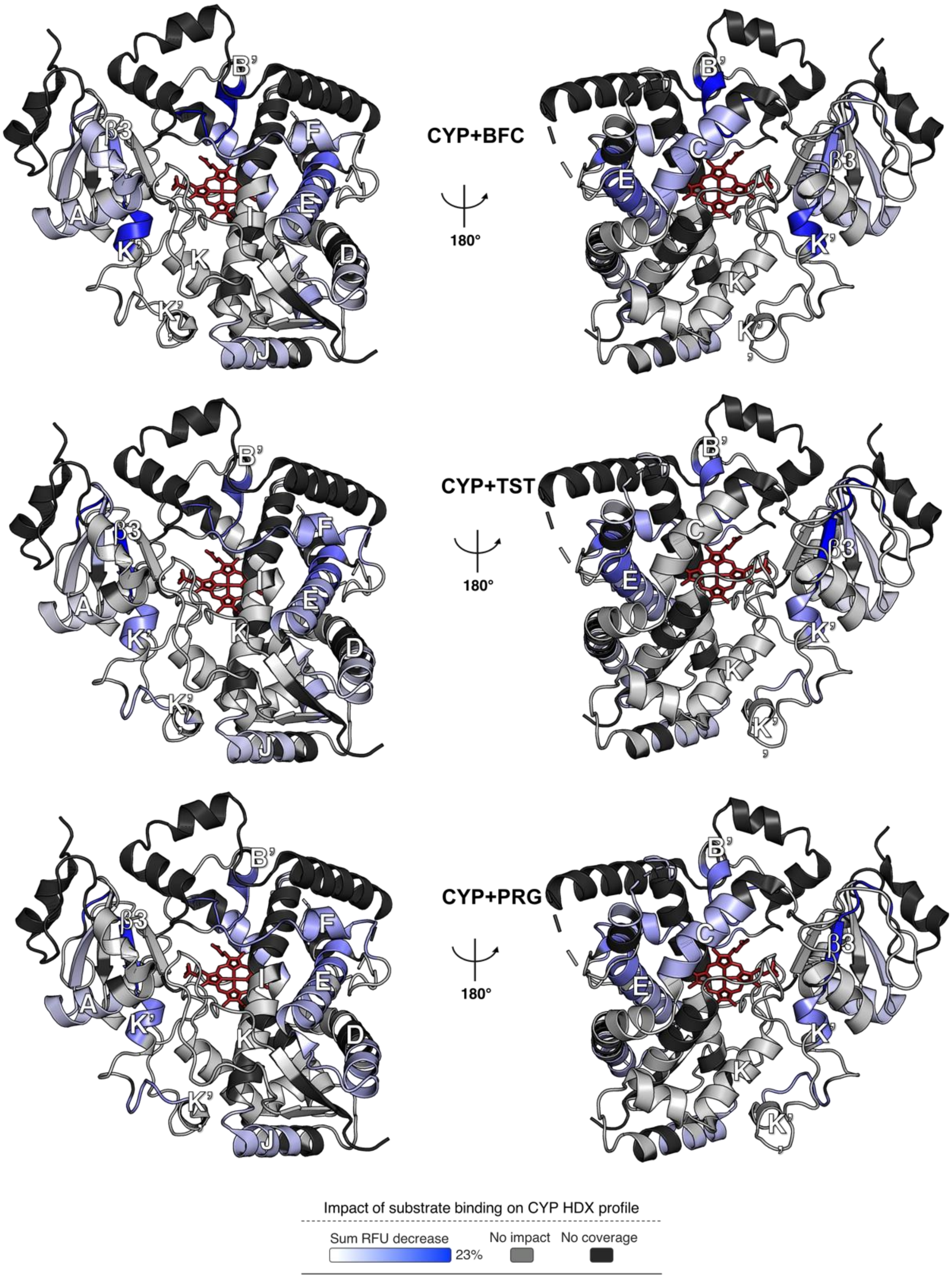


**Figure S9.** Impact of substrate binding on the structural dynamics of CYP3A4. Differential HDX profiles comparing CYP to either CYP+BFC, CYP+TST or CYP+PRG are shown. The significant sum RFU difference (CI 99%) of two time points (5, 60 min) is mapped onto the CYP3A4 structure (PDB: 1W0F), shown in both distal (left) and proximal (right) views. The blue color indicates a segment undergoing a decrease in deuterium uptake upon interaction with the substrate. The light gray color reveals regions unaffected by substrate binding, whereas black is used to show regions non-covered in the MS analysis.

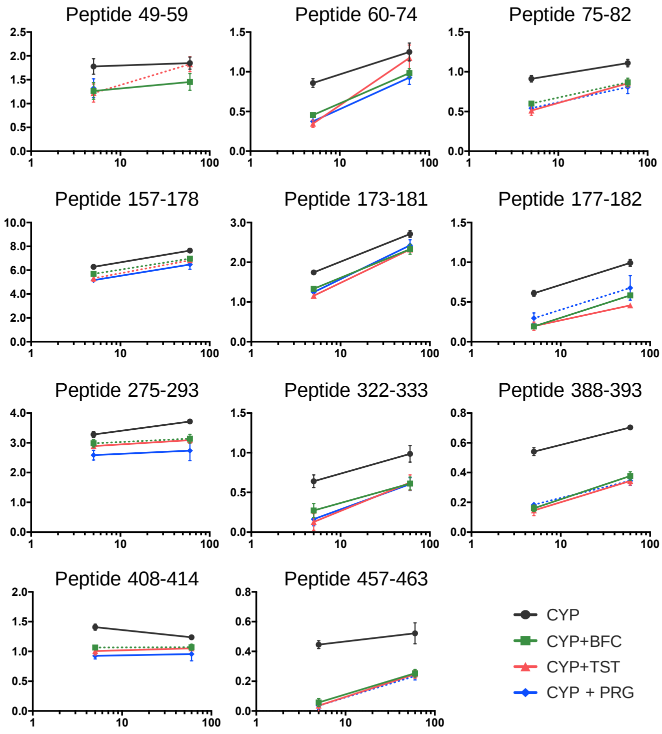


**Figure S10.** Deuterium uptake plots for all regions showing a significant change in the presence of substrate. The y-axis represents the deuterium uptake (Da) and the x-axis is time in a logarithmic scale. Black curves are for CYP alone; green, pink and blue curves are for CYP in the presence of BFC, TST and PRG, respectively. Dashed curves indicate that the difference in deuterium uptake between CYP was not significant (CI 99%).
